# Addiction-associated genetic variants implicate brain cell type- and region-specific cis-regulatory elements in addiction neurobiology

**DOI:** 10.1101/2020.09.29.318329

**Authors:** Chaitanya Srinivasan, BaDoi N. Phan, Alyssa J. Lawler, Easwaran Ramamurthy, Michael Kleyman, Ashley R. Brown, Irene M. Kaplow, Morgan E. Wirthlin, Andreas R. Pfenning

## Abstract

Recent large genome-wide association studies (GWAS) have identified multiple confident risk loci linked to addiction-associated behavioral traits. Genetic variants linked to addiction-associated traits lie largely in non-coding regions of the genome, likely disrupting cis-regulatory element (CRE) function. CREs tend to be highly cell type-specific and may contribute to the functional development of the neural circuits underlying addiction. Yet, a systematic approach for predicting the impact of risk variants on the CREs of specific cell populations is lacking. To dissect the cell types and brain regions underlying addiction-associated traits, we applied LD score regression to compare GWAS to genomic regions collected from human and mouse assays for open chromatin, which is associated with CRE activity. We found enrichment of addiction-associated variants in putative **CREs** marked by open chromatin in neuronal (NeuN+) nuclei collected from multiple prefrontal cortical areas and striatal regions known to play major roles in reward and addiction. To further dissect the cell type-specific basis of addiction-associated traits, we also identified enrichments in human orthologs of open chromatin regions of mouse **neuronal subtypes: cortical excitatory, D1, D2, and PV**. Lastly, we developed machine learning models from mouse cell type-specific regions of open chromatin to further dissect human NeuN+ open chromatin regions into cortical excitatory or striatal D1 and D2 neurons and predict the functional impact of addiction-associated genetic variants. Our results suggest that different **neuronal subtypes** within the reward system play distinct roles in the variety of traits that contribute to addiction.

**Significance Statement:** **We combine statistical genetic and machine learning techniques to find that the predisposition to for nicotine, alcohol, and cannabis use behaviors can be partially explained by genetic variants in conserved regulatory elements within specific brain regions and neuronal subtypes of the reward system.** This computational framework can flexibly integrate **open chromatin** data across species to screen for putative causal variants in a cell type-and tissue-specific manner across numerous complex traits.

## INTRODUCTION

Substance use disorders (SUD) have increased in prevalence over the last three decades, with an estimated 100 million cases worldwide (GBD 2016 Alcohol and Drug Use Collaborators, 2018; Eddie et al., 2019). Pharmacological interventions are limited in their ability to cure addiction due to physiological and logistical barriers (Pullen and Oser, 2014; Pear et al., 2019). As the societal epidemic of substance use grows, there is a greater need to understand the neurobiology of substance use behaviors and addiction.

The reward circuits co-opted in addiction **as well as the associated neural cell types** are highly conserved across primates and rodents **(Monaco et al., 2015; Grillner and Robertson, 2016; Scaplen and Kaun, 2016; Hodge et al., 2019)**. It is generally accepted that addictive substances promote impulsive and compulsive behavior by activating the mesolimbic dopamine system, in which dopaminergic inputs from the ventral tegmental area project to medium spiny neurons (MSN) of the nucleus accumbens (NAc) **in the ventral striatum (STR)** (Koob and Volkow, 2010). Glutamatergic inputs to the NAc from the amygdala, frontal cortex, and hippocampus contribute to motivational action through the extrapyramidal motor system (Koob and Volkow, 2010). Subsequently, the NAc sends outputs to nuclei of the ventral pallidum**, which are** critical for processing and modulating substance reward signal (Koob and Volkow, 2010). **The development of compulsive substance-seeking is hypothesized to be linked to recruitment of the dorsal STR, which together with the prefrontal cortical regions regulates a variety of reward and addiction-related phenotypes** (Koob and Volkow, 2010; Goldstein and Volkow, 2011). **These** findings emphasize **that substance abuse behavior involves** the interplay of **the** brain regions and **cell types that make up the reward system.**

Increasing evidence reveals strong genetic links to substance use risk (Pasman et al., 2018; Erzurumluoglu et al., 2019; Karlsson Linnér et al., 2019; Liu et al., 2019b) and SUD (Kendler and Prescott, 1998a, 1998b; Dick, 2016; Waaktaar et al., 2018). Genome-wide association studies (GWAS) report that genetic risk for substance use shares underlying architecture with other neuropsychiatric disorders (Pasman et al., 2018; Liu et al., 2019b), of which risk variants tend to lie in non-coding, functional regions of the human genome (Jensen, 2016). These **genetic variants, including** single nucleotide polymorphisms (SNPs), can disrupt transcription factor binding in cis-regulatory elements (CREs) with varying impact on gene regulation and downstream neural circuitry. Many CREs have tissue- and cell type-specific activity (Roadmap Epigenomics Consortium et al., 2015), suggesting that cell types and tissues underlying addiction may be uniquely targeted by genetic variants at these CREs. GWAS for nicotine-, alcohol- (Liu et al., 2019b), and cannabis-use traits (Pasman et al., 2018) have identified multiple confident risk loci and SNPs linked to addiction-associated phenotypes with brain-specificity, yet their effects on the CREs of specific brain regions and cell types involved in addiction pathophysiology are an open area of inquiry.

**A comparison of GWAS to functional annotations of the human genome** have **yielded estimates that over 90% of SNPs associated with complex phenotypes lie** within functional non-coding regions**, which are marked by epigenetic features including open chromatin. (Maurano et al., 2012; Finucane et al., 2015)**. Linkage disequilibrium (LD) of significant SNPs complicates the identification of causal variants contributing to genetic risk (Bush and Moore, 2012). **Regression of SNP LD scores against GWAS summary statistics (LDSC regression) is the dominant method for relating human genetics to functional annotations. LDSC regression partitions risk SNPs identified by GWAS into the tissues or cell types in which genetic variation in CREs may contribute to heritability of complex traits (Finucane et al., 2015; Visscher et al., 2017)**. **Yet**, the functional consequences of risk SNPs in CRE sequences cannot be reliably inferred from DNA sequences alone (Shlyueva et al., 2014). Recent developments in epigenomic assays (Buenrostro et al., 2013; Mo et al., 2015; Tak and Farnham, 2015) and machine learning (Ghandi et al., 2014; Zhou and Troyanskaya, 2015; Kelley et al., 2016, 2018; Lee, 2016) can predict cell types affected by addiction-associated genetic variation to propose cell type-specific hypotheses on the pathogenesis of addiction.

Here, we implement a framework that **links the genetic predisposition to addiction-associated traits to specific brain** regions and cell types **within them by identifying which have open chromatin regions that are enriched for SNPs identified by GWAS**. We first **intersect SNPs measured by GWAS** across human and mouse bulk tissue and cell type-specific open chromatin **regions** to identify **putative** region- and cell type-specific CREs that may be impacted by genetic variation associated with addiction-related traits. **To overcome limits of cellular resolution in the human brain, we apply convolutional neural network models trained on transgenically-labelled neuron populations in the reward system of mice to predict the cell type-specificity of GWAS-associated SNPs in the human genome. We further** apply these models to **the problem of** screen**ing** for putative causal SNPs within dense loci reported in GWAS for addiction-associated traits. This pipeline, to our knowledge, describes the first integrative analyses across species, **brain** regions and cell types to screen for candidate causal addiction-associated genetic risk variants in dense loci with numerous significant SNPs in LD.

## RESULTS

**Genetic risk for substance use traits is associated with the neuronal epigenomes of reward areas** Recent well-powered GWAS have **identified dozens of candidate genetic risk loci associated with seven addiction-associated traits: age of smoking initiation (AgeOfInitiation), average number of cigarettes smoked per day (CigarettesPerDay), having ever regularly smoked (SmokingInitiation), being a former versus current smoker (SmokingCessation), the number of alcoholic drinks per week (DrinksPerWeek), and lifetime cannabis use (Cannabis), and risk tolerance (RiskyBehavior)** (Pasman et al., 2018; Karlsson Linnér et al., 2019; Liu et al., 2019b). These GWAS **measure** reward, risk tolerance, and various substance use behaviors**, thereby providing** a means of studying genetic variation associated with addiction. We found that 72-98% of addiction-associated genetic variants **lie in** non-coding regions of the genome (**Figure 1A**). **Of those risk variants, 47-85% lie in introns, which is a substantial over-representation in each GWAS** (odds ratio, OR_AgeOfInitiation_ =2.3, OR_Cannabis_ = 2.3, OR_CigarettesPerDay_ = 1.4, OR_DrinksPerWeek_ =1.6, OR_RiskyBehavior_ = 1.4, OR_SmokingCessation_ = 1.8, OR_SmokingInitiation_ =1.3, Fisher’s Exact P_Bonferroni_ < 8 x 10^-79^). Furthermore, **pairwise genetic correlations of risk alleles in these seven GWAS indicated shared and distinct genetic architecture across addiction-associated traits** (r_g_**,Supplemental Figure 1A**). Although common genetic variants are shared between addiction-associated traits on a genome-wide scale, the reported significant loci are often unique to a particular trait and are densely packed with SNPs in high LD **(Supplemental Figure 1B**). **SNPs that are associated with the seven traits span 205 non-overlapping loci across the human genome and include on average 71 SNPs (minimum 1, median 22.5, maximum 1780) within**each locus that are either genome-wide significant (P_GWAS_ < 5 x 10^-8^) or in high LD with the lead SNPs (R^2^ > 0.8, Supplemental Table 1).

**Figure 1.**
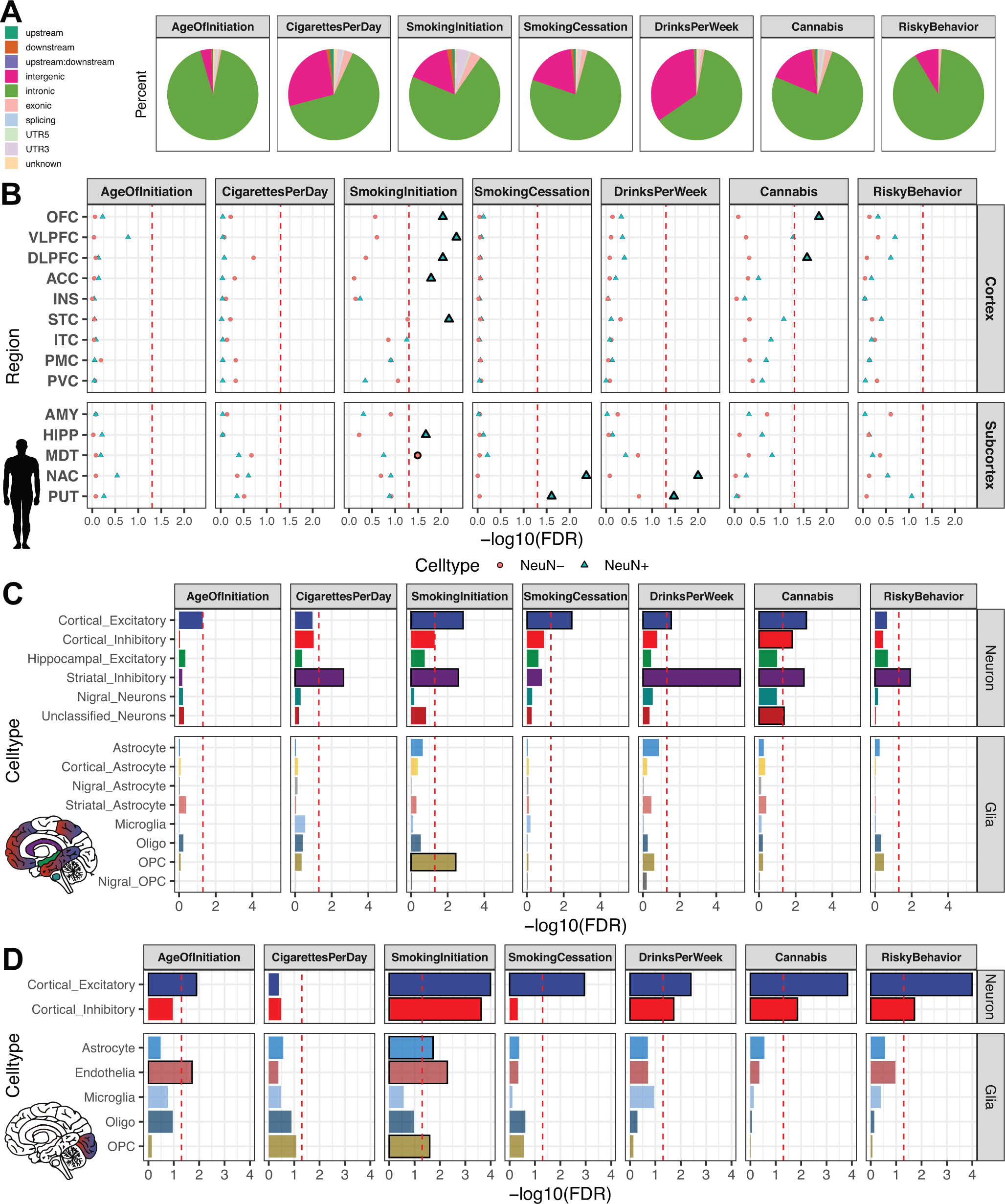
Substance use and risky behavior GWAS risk variants enrich within reward region- and cell type-specific epigenomic profiles. Partitioned LDSC regression (GWAS enrichment) finds enrichment of substance use and risky behavior traits in region-specific and cell type-specific open chromatin profiles of human postmortem brain. **(A)** Pie chart of ANNOVAR-annotated (Wang et al., 2010) SNP function of addiction-associated trait lead and off-lead SNPs in LD R^2^ > 0.8. Dark colors indicate un-transcribed/non-coding annotations, light for transcribed/exonic annotations. SNP annotation labels are according to ANNOVAR using ENSEMBL build 85 gene annotations (Methods). **(B)** GWAS enrichment false-discovery rates in ATAC-seq of 14 postmortem human brain regions coupled with NeuN-labeled fluorescence activated nuclei sorting (Fullard et al., 2018). Brain regions are stratified by cortical and subcortical regions, with cortical regions ordered frontal to caudal. Sorted cell types within each brain region are denoted by shape (blue triangle for NeuN+/neuronal, red circle for NeuN-/glial). FDR-adjustment was performed across all enrichments on the Fullard *et al*. dataset for Figure 1B and Figure 3: **Cell type-specific convolutional neural network (CNN) models refine human NeuN+ enrichments for substance use genetic risk GWAS. (A**) **Schematic** to predict cell type-specific activity of NeuN+ ATAC-seq peaks enriched from brain regions assayed in Fullard *et al*. (Fullard et al., 2018) using **CNN** models trained on mouse cell-type specific ATAC-seq peaks. **CNN**-predicted OCRs are input into GWAS enrichment. (**B)** Partitioned LD score regression of addiction associated traits in Fullard *et al*. NeuN+ OCRs predicted to be cell type-specific by machine learning models of open chromatin. Cell types are colored by the source mouse cell type-specific OCRs from **Error! Reference source not found.A**. Original enrichments from Figure 1A are reproduced in black. **Larger, bolded points are significant for FDR < 0.05 (red dotted line)**.

We investigated whether genetic variants **implicated by** addiction-associated GWAS **show a tendency to** cluster at putative cis-regulatory elements (CREs) of the brain using a partitioned heritability LDSC regression approach**, which looks for an enrichments of significant SNPs from GWAS in human annotations** (Bulik-Sullivan et al., 2015b; Finucane et al., 2018). **We applied LDSC to compare the seven addiction-associated GWAS to** open chromatin region (OCR) annotations of sorted neuronal (NeuN+) and glial (NeuN-) nuclei across 14 brain regions(Fullard et al., 2018) (**Figure 1B**). We found that genetic variants associated with SmokingInitiation, SmokingCessation, DrinksPerWeek, and Cannabis significantly enriched in NeuN+ OCRs of brain regions known and speculated to contribute to reward and addiction(Volkow and Morales, 2015) (FDR < 0.05). **We** found that genetic variants associated with SmokingInitiation and Cannabis shared enrichment in NeuN+ prefrontal cortical OCRs (from orbitofrontal cortex and dorsolateral prefrontal cortex) while those associated with SmokingCessation and DrinksPerWeek shared enrichment in NeuN+ striatal OCRs (both putamen and NAc). The enrichments of NeuN+ OCRs indicate that genetic variation in epigenomes of neuronal populations from frontal cortex and striatum contribute to addiction liability. The difference in NeuN+ enrichments between regions across addiction-associated traits can likely be explained by the difference in proportions **and identities** of neuronal subtypes of each area, so we sought to dissect the different neuronal subtypes contributing to these enrichments.

Broad marker-gene based labeling approaches, such as using NeuN to label neurons, do not capture the rich diversity of neuronal subtypes; bulk NeuN+ open chromatin signal represents an average signal from heterogeneous neuronal subtypes, each with distinct epigenomic landscapes, gene regulation, network connectivity. Hence, NeuN-labeled open chromatin profiles likely do not capture OCRs unique to less populous neuronal subtypes. The difference in proportions of neuronal subtypes between brain regions may also contribute to brain region-specific NeuN+ OCR enrichment for GWAS variants of addiction-associated traits. We therefore applied LDSC regression GWAS enrichment on **single cell** open chromatin profiles from human postmortem **isocortical, striatal, hippocampal, nigral (Figure 1C) and occipital cortical cell types** (Lake et al., 2018; Corces et al., 2020) (**Figure 1D**). We found that addiction-associated genetic variants largely enriched in both excitatory and inhibitory neuronal OCRs. **Genetic variants associated with SmokingInitiation, SmokingCessation, DrinksPerWeek, and Cannabis enriched in isocortical excitatory neuron OCRs (Figure 1C). We found enrichment of genetic variants associated with CigarettesPerDay, SmokingInitiation, SmokingCessation, DrinksPerWeek, Cannabis, and RiskyBehavior in striatal inhibitory neurons. Genetic variants associated with Cannabis also enriched in isocortical inhibitory neuron and unclassified neuron OCRs. Among the glial cell types, only oligodendrocyte precursor cell OCRs were enriched for an addiction-associated trait (SmokingInitiation).** We found enrichment of genetic variants associated with AgeOfInitiation and SmokingCessation in OCRs of **occipital** cortical excitatory neurons. We found no enrichment of genetic variants associated with CigarettesPerDay for OCRs of occipital cortex cell types. Genetic variants associated with SmokingInitiation, which enriched in astrocyte, endothelial, inhibitory, and oligodendrocyte precursor cell OCRs **from occipital cortex**, shared enrichment in NeuN-OCRs of mediodorsal thalamus (**Figure 1B**). Interestingly, genetic variants associated with SmokingCessation, which **showed enrichment** for striatal NeuN+ OCRs, enriched only for OCRs of **occipital** cortical excitatory neurons and not cortical inhibitory neurons. Sorted bulk ATAC-seq only showed enrichment of SmokingCessation associated genetic variants in OCRs of NeuN+ striatal regions, which are largely composed of inhibitory MSNs. **We overall found that the enrichments of addiction-associated genetic variants in Corces *et al*. isocortex OCRs agreed with those in Lake *et al*. occipital cortex OCRs**. Single-cell epigenomics of human postmortem brain can further dissect the genetic risk for substance-use traits into neuronal subtypes that otherwise would not be parsed with bulk tissue assays.

We confirmed that our pipeline for LDSC regression on NeuN-sorted OCRs from 14 brain regions is able to reproduce the GWAS enrichments published by Fullard *et al*. While our approach uses OCRs from reproducible ATAC-seq peaks rather than differentially accessible peaks, we found consistent enrichments of genetic variants associated with schizophrenia risk (Schizophrenia), highest level of educational attainment (EduAttain), and habitual sleep duration (SleepDuration) **(Supplemental Figure 2B)**. We did not find enrichment in brain OCRs of genetic variants identified in several low-powered GWAS (cocaine dependence (CocaineDep) (Cabana-Domínguez et al., 2019), opioid dependence (OpioidDep) (Cheng et al., 2018), and obsessive-compulsive disorder (OCD) (International Obsessive Compulsive Disorder Foundation Genetics Collaborative (IOCDF-GC) and OCD Collaborative Genetics Association Studies (OCGAS), 2018), each of which had included fewer than 5000 individuals with the trait **(Supplemental Figure 2A)**. In addition, we found no enrichments in brain OCR for several well-powered studies of traits related to addiction behaviors, **including** multi-site chronic pain (ChronicPain) (Johnston et al., 2019) and cups of coffee per day (CoffeePerDay) (Coffee and Caffeine Genetics Consortium et al., 2015). We also found no enrichment in brain OCRs for anthropometric traits, **including** coronary artery disease (CAD) (Howson et al., 2017), bone mineral density (BMD) (Kemp et al., 2017), and lean body mass (LBM) (Zillikens et al., 2017) **(Supplemental Figure 2B, C)**. Lastly, we validated that human OCRs from non-brain tissues would not enrich for risk variants associated with brain traits. We gathered publicly available OCRs from stomach ATAC-seq, adipocyte ATAC-seq, preadipocyte ATAC-seq, liver DNase-seq, and lung DNase-seq profiles (ENCODE Project Consortium, 2012; Thurman et al., 2012; Davis et al., 2018; Cannon et al., 2019) **(Supplemental Figure 4D)** and performed LDSC regression on the total 18 GWAS from above. To our expectation, we did not find enrichments of stomach, liver, or lung OCRs for genetic variants associated with brain-related traits. We did find enrichment of BMD in lung OCRs, a connection previously recognized (Lee et al., 2016; Kim et al., 2019; Zeng et al., 2019). The secondary GWAS enrichments in other traits and foregrounds demonstrate two trends: a GWAS trait would enrich if the GWAS was properly powered to detect genetic risk variants, and the foreground regions are from cell types or tissue of that trait’s potential etiological origin.

### Mouse-human conserved cell type-specific open chromatin enrich for addiction risk loci

In order to **further interrogate the** different neuronal subtypes that comprise the enrichment of addiction-associated genetic variants in OCR sets measured by Fullard *et al*., Lake *et al*., and Corces *et al*. (Figure 1, Supplemental Figure 2), we performed targeted epigenomic **experiments** in mouse **on isolated neuronal subtypes** from **key brain regions of the reward circuit:** frontal cortex (CTX), caudoputamen (CPU), and the nucleus accumbens (NAc). We isolated **nuclei from specific cell types** for ATAC-seq using a modified version of the INTACT approach (Mo et al., 2015) called *<u>c</u>re*-<u>s</u>pecific <u>n</u>uclei <u>a</u>nchored independent labeling (cSNAIL). cSNAIL-INTACT **was applied** to isolate nuclei marked by *Pvalb*, *Sst*, *Drd1*, and *Adora2a* in *cre-driver* lines using a shortened form of the *Sun1-Gfp* fusion protein packaged with AAV-PHP.eb and delivered through retro-orbital injection (Figure 2A). We show that cell type-targeting provided markedly distinct genome-wide ATAC-seq profiles compared to bulk tissue ATAC-seq alone (**Supplemental Figure 3A**). cSNAIL ATAC-seq specifically captured nuclei with increased accessibility around the marker gene that was driving *Cre* recombinase expression (**Supplemental Figure 3B**). Accessibility around cSNAIL ATAC-seq transcription start sites (TSS) strongly correlated with matched pseudobulk gene expression in the same cell type and tissue (**Methods**, both Pearson and Spearman correlation P_bonf_ < 2 x 10^-16^ **Supplemental Figure 3C,D**). We applied HALPER**, an approach that leverages reference-free multi-species genome alignments to produce 1-1 contiguous CRE orthologs** (Zhang et al., 2020), to reliably map ∼70% of mouse **neuronal subtype** OCRs to their human orthologs in **the hg38 human reference genome (Methods) for LDSC regression GWAS analysis**.

**Figure 2:**
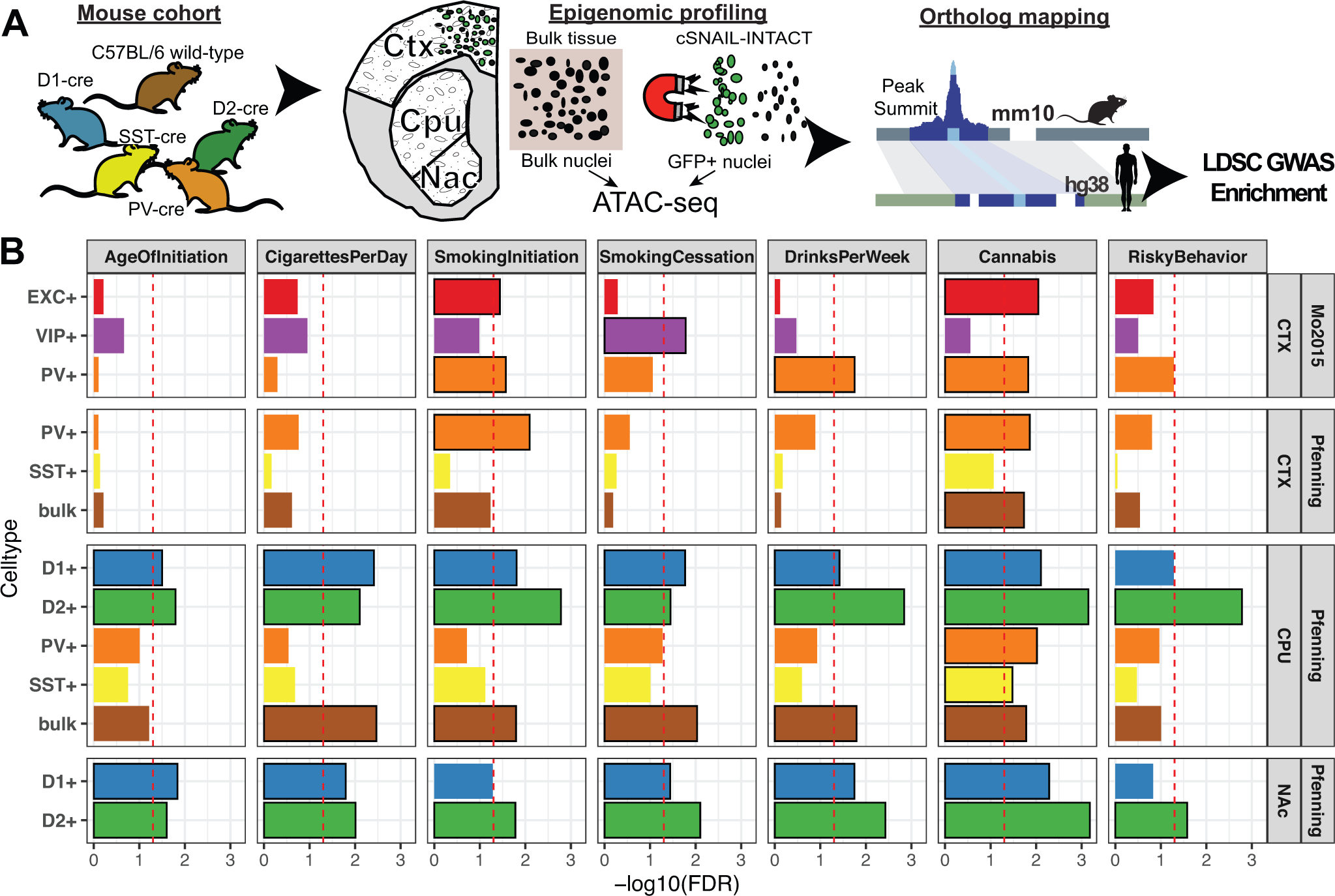
Cell type-specific enrichment of substance use traits are conserved in mouse-human orthologous open chromatin regions. **(A)** Experimental design to map human orthologous regions from mouse ATAC-seq of bulk cortex (CTX), dorsal striatum (CPU), and nucleus accumbens (NAc) of cre-dependent Sun1-GFP Nuclear Anchored Independent Labeled (cSNAIL) nuclei of D1-cre, D2-cre, PV-cre, and SST-cre mice. cSNAIL ATAC-seq experiments report enriched (+) nuclei populations. **(B)** Partitioned LD score regression finds enrichment of substance use and risky behavior traits for brain region and cell type specific ATAC-seq open chromatin profiles of mouse brain. Replication of enrichment is shown using INTACT-enriched OCRs from Mo *et al* (Mo et al., 2015) of cortical excitatory (EXC+), vasoactive intestinal peptide interneuron (VIP+), and parvalbumin interneuron (PV+). Enrichments that are enriched at FDR < 0.05 are plotted with black outlines. FDR-adjusted p-value was performed across all mouse-human ortholog GWAS enrichment across Figure 2.

Our GWAS enrichment **analysis** of human orthologs from mouse OCRs **(mouse-human orthologs) measured in** various neuronal subtypes and bulk tissue (**Figure 2B**) show that genetic variants associated with SmokingInitiation and Cannabis shared enrichment in cortical PV and EXC neuron OCRs from both Mo *et al*. and this study (**Pfenning data,** FDR < 0.05). **Genetic variants associated with Cannabis** further enriched in CTX bulk tissue OCRs, which could be attributed to signal from cortical EXC and PV neuron populations. Cortical PV neuron OCRs further enriched with genetic variants associated with DrinksPerWeek. SmokingCessation associated genetic variants distinctly enriched in cortical VIP neuron OCRs.

Within **neuronal subtypes** from CPU and NAc, we found enrichment of genetic variants associated with all measured addiction-associated traits in CPU and NAc D2 MSN OCRs. Genetic variants associated with all measured traits excluding SmokingInitiation and RiskyBehavior all enriched in CPU and NAc D1 MSN OCRs. CPU D1 MSN OCRs were enriched with genetic variants associated with all measured traits excluding RiskyBehavior. We found that CPU bulk tissue OCRs were enriched with genetic variants associated with all measured addiction-associated traits excluding AgeOfInitiation and RiskyBehavior. Distinctly, CPU PV+ and SST+ neuron OCRs enriched with genetic variants associated with Cannabis.

Corresponding to our analysis of human brain OCRs, we also confirmed the specificity of mouse-human orthologous CRE enrichments for genetic variants **associated with** addiction-related, brain-related, and non-brain related traits **(Supplemental Figure 4)**. We found enrichments of genetic variants associated with ChronicPain in cortical PV neuron OCRs from both Mo *et al*. and this study **(Supplemental Figure 4A).** Within striatal cell types, we found that CPU D2 and NAc D1 MSN OCRs were enriched for genetic variants associated with ChronicPain. In contrast, CPU D1 and NAc D2 MSN OCRs were enriched for genetic variants associated with OpioidDep. Genetic variants associated with OpioidDep also enriched in CPU D1 MSN and CPU PV OCRs. Schizophrenia, EduAttain, and SleepDuration associated genetic variants all enriched in OCRs of all measured cell types **(Supplemental Figure 4B)**. None of these **mouse-**human orthologs enriched for genetic variants associated with non-brain-related traits: BMD, CAD, and LBM **(Supplemental Figure 4C)**. We validated that our approach to map OCRs from mouse to human did not bias enrichment to brain traits by performing GWAS enrichment on OCRs from mouse non-brain tissues (kidney, liver, and lung) (**Supplemental Figure 4D)**. As expected, we did not find an enrichment for genetic variants associated with a brain-related trait. We did find that **mouse-**human orthologs of lung OCRs enrich for BMD, which concords with the enrichment of human lung OCRs.

### Convolutional Neural Network (CNN) models of mouse cell type-specific CRE activity refine human NeuN+ OCRs for GWAS enrichment

The genetic tools available for mouse research allowed us to isolate the nuclei of specific neuronal subtypes and generate deep open chromatin profiles at greater cellular resolution. However, a lack of mouse-human conservation in the cell type-specificity of CREs could lead to false negatives and false positives at specific loci that add noise to GWAS comparisons. To leverage the strengths of the mouse and human approaches, we developed a procedure to predict the neuronal subtype-specificity of human OCRs using machine learning models trained in mouse. The OCR profile of each neuronal subtype is a result of a developmental cascade of transcription factors that cooperatively recognize and bind to specific sequence elements in the genome, resulting in a neuronal subtype-specific open chromatin profile (Spitz and Furlong, 2012). These complex combinations of sequence features comprise regulatory code that links genome sequence to neuronal subtype-specific open chromatin. This regulatory code can be effectively learned using convolutional neural networks (CNNs) and has been demonstrated to be highly conserved between mouse and human (Zhou and Troyanskaya, 2015; Chen et al., 2018)

The concordant pattern of enrichment for addiction associated genetic variants in human and mouse-human orthologous OCRs suggested that risk variants may affect the regulatory **activity of neuronal subtypes** conserved **between human and** mouse. We therefore **devised and trained a collection of CNN binary classification models to learn the genome sequence features that distinguish OCRs** for cortical excitatory (EXC) neurons, striatal D1 MSNs, and striatal D2 MSNs (Zhou and Troyanskaya, 2015; Kelley et al., 2016, 2018; Chen et al., 2018). For each set of reproducible OCRs from mouse INTACT and cSNAIL group, we trained 5-fold cross-validated models to predict the reproducible peaks from ten times the number of **nucleotide content**-matched negative sequences **(Methods)**. Our models made confident predictions on held-out test sequences as reported by **high F1-scores, high area under the precision-recall curves (Supplemental Figure 5A)**, **and low false positive rates at a blind threshold of 0.5 (Supplemental Figure 5B). These models reproducibly learned transcription factor motif families that are enriched in human neuronal subtypes of cortex (MEF2, JUN) and striatum (POU, NRF1, ZFHX3), as previously reported by Fullard *et al*. (Supplemental Figure 5F, Supplemental Table 2)**.

We reasoned that NeuN+ OCR signal**, which is comprised of OCR signals from several neuronal subtypes, can be parsed into its component cell types by CNNs that are trained to predict OCR activity in those component cell types**. **This enables the study of human addiction genetics at a cell type-level resolution from high-quality tissue-level open chromatin profiles**. To discern whether NeuN+ OCR enrichments in addiction-associated genetic variants come from the same cell types **observed** in **Figure 3**, we **applied our trained CNN** models to predict whether **bulk** cortical or striatal NeuN+ OCRs have activity in **either** cortical EXC or striatal D1 and D2 **neurons**, respectively (**Figure 3A**). We did not conduct these analyses for PV neurons because they comprise a much lower percentage of cortical and striatal neurons than the other neuron types (Beaulieu, 1993; Lefort et al., 2009). We ran LDSC regression (Finucane et al., 2018) GWAS enrichments on the sets of NeuN+ OCRs predicted to be specific to cortical EXC, striatal D1, and striatal D2 neurons. Genetic variants associated with SmokingInitiation, which initially enriched in OCRs of various NeuN+ frontal cortical areas (**Figure 1B**), enriched in NeuN+ OCRs predicted to be active in EXC neurons (**Figure 3B**). Genetic variants associated with Cannabis, which enriched in NeuN+ cortical OCRs (**Figure 1B**), also enriched in NeuN+ OCRs predicted to be active in EXC neurons. The enrichments of excitatory cortical cell type-specific OCRs for SmokingInitiation and Cannabis associated genetic variants agree with the results from the **analysis of the** Fullard *et al*., Corces et al., and Lake *et al*. **OCR** datasets (**Figure 1B, C)**. Genetic variants associated with SmokingCessation and DrinksPerWeek, which enriched in PUT and NAc NeuN+ OCRs (**Figure 1B**), shared enrichment in OCRs predicted active in both D1 and D2 MSNs of both PUT and NAc. The framework that we outline in **Figure 3A** refines addiction genetic risk signal to neuronal subtypes **and** maintains the brain region context of the source NeuN+ OCR. **This framework can be applied to CREs from any tissue-cell type combination for which bulk tissue open chromatin measurements are available from human and cell type open chromatin measurements are available from another vertebrate** (Chen et al., 2018; Minnoye et al., 2020).

**Figure 3:**
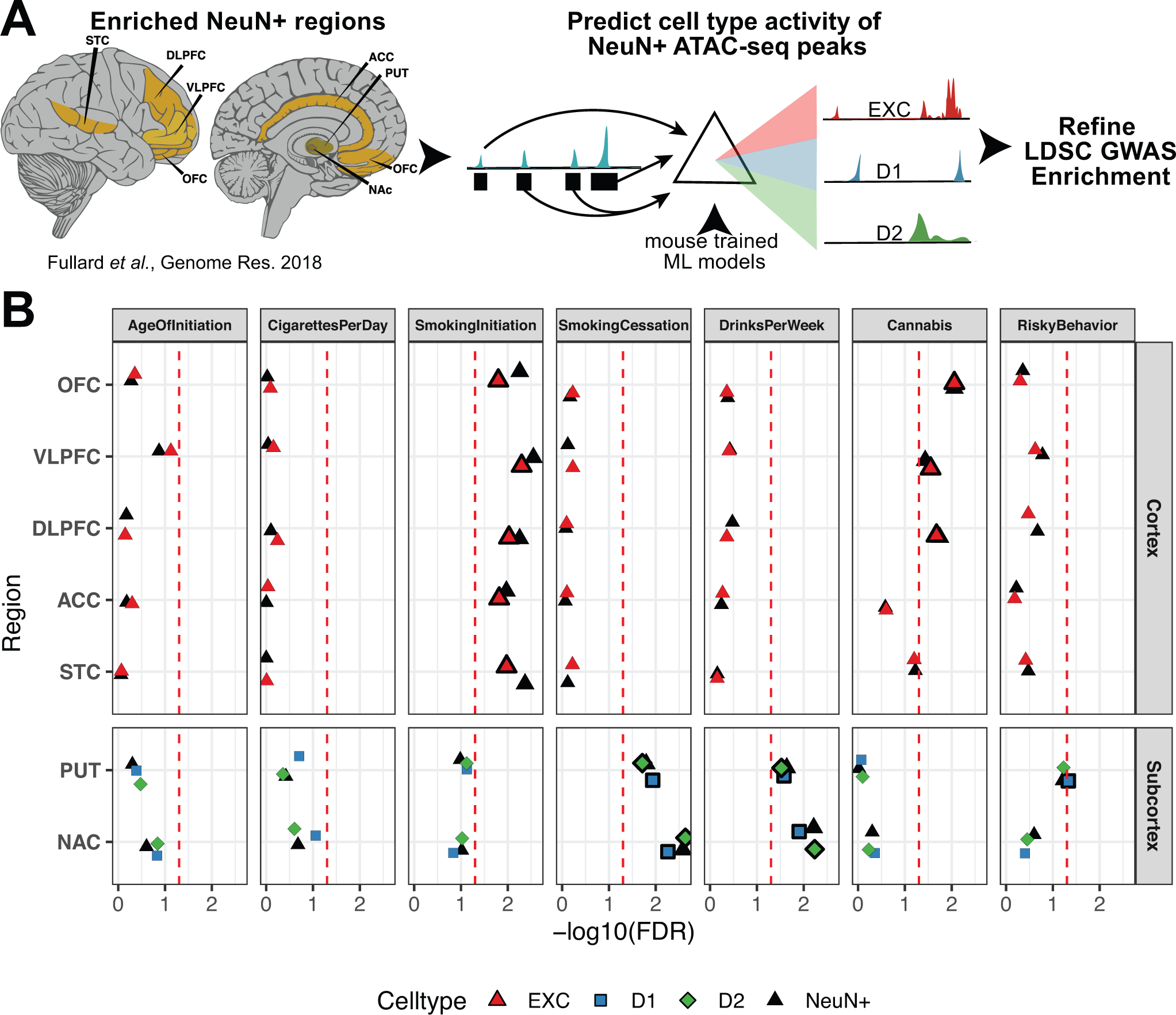
Cell type-specific convolutional neural network (CNN) models refine human NeuN+ enrichments for substance use genetic risk GWAS. **(A**) **Schematic** to predict cell type-specific activity of NeuN+ ATAC-seq peaks enriched from brain regions assayed in Fullard *et al*. (Fullard et al., 2018) using **CNN** models trained on mouse cell-type specific ATAC-seq peaks. **CNN**-predicted OCRs are input into GWAS enrichment. (**B)** Partitioned LD score regression of addiction associated traits in Fullard *et al*. NeuN+ OCRs predicted to be cell type-specific by machine learning models of open chromatin. Cell types are colored by the source mouse cell type-specific OCRs from **Error! Reference source not found.A**. Original enrichments from Figure 1A are reproduced in black. **Larger, bolded points are significant for FDR < 0.05 (red dotted line)**.

### Convolutional Neural Network (CNN) models predict allele-specific activity of addiction-associated GWAS SNPs in neuronal subtypes

Lastly, we applied our convolutional neural network (**CNN)** models to screen addiction-associated genetic variants for predicted functional activity in EXC, D1, and D2 neuronal subtypes. **CNN**-based approaches have been demonstrated to fine-map dense risk loci and select candidate causal genetic variants (Alipanahi et al., 2015; Zhou and Troyanskaya, 2015; Kelley et **al., 2016, 2018; Corces et al., 2020), yet none have been applied in the context of addiction-associated genetic risk or in the cell types that we have assayed.** We identified 14,790 unique SNPs that were collected across the seven addiction-associated GWAS to score for differential neuronal subtype OCR activity (Methods). We expect that many SNPs reported from GWAS are significantly associated with traits due to LD rather than being the true causal variant. When scored with our CNN models, the 96.2% of addiction-associated SNPs that either do not lie in any OCR or in only NeuN-OCR have low probability to be active in excitatory, D1, or D2 neuronal subtypes. We also found that these SNPs have significantly lower predicted probability of activity than the remaining 3.8% of addiction-associated SNPs in any NeuN+ OCR (P_Bonferroni_ < 0.05, Figure 4A). We then predicted the probability of activity for both the effect and non-effect allele and estimated the differential impact of the alleles in order to fine-map candidate causal effect SNP and target neuronal subtype and tissue. Most SNPs do not have predicted differential allelic activity in a neuronal subtype, while a handful of SNPs have larger differential activity that deviate from a normal distribution when visualized on quantile-quantile plots (Supplemental Figure 5C, Methods). We outline in Supplemental Figure 5D an approach to prioritize the candidate causal SNPs by two lines of evidence: 1) a predicted differential neuronal subtype OCR activity with large effect size that is controlled for false discovery (q-value < 0.05, Methods) and 2) having physical overlap with measured human NeuN+ OCR in Fullard *et al*. (Supplemental Figure 5D). We are able to prioritize 55 SNPs spanning 37 loci to Tier A which have both significant predicted ΔSNP probability effect and overlaps a Fullard *et al*. NeuN+ OCR, 505 SNPs to Tier B that only have predicted ΔSNP probability **effect, and 502 SNPs to Tier C as overlapping NeuN+ open chromatin without a predicted significant ΔSNP probability effect (Supplemental Table 1).**

**Figure 4:**
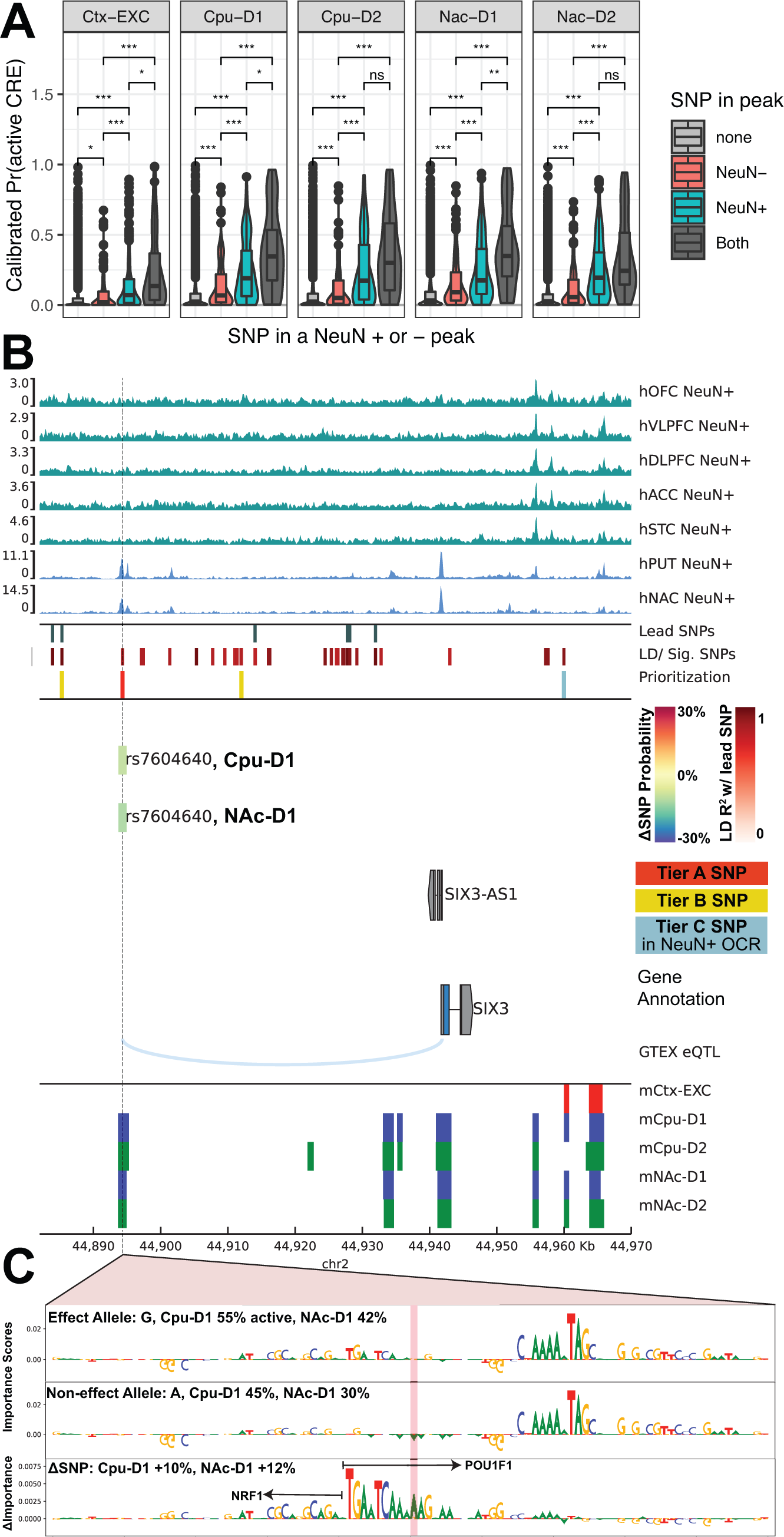
Convolutional Neural Network (CNN) models for predicting cell type-specific open chromatin predict activity of addiction GWAS SNPs. **(A) Cell type-activity predicted probability active by each set of CNN models of cell type activity for genome-wide significant SNPs and off-lead SNPs in LD R^2^ > 0.8 with the lead SNPs.** Activity scores for SNPs are stratified by overlap with Fullard *et al*. (Fullard et al., 2018) cortical or striatal **NeuN+ (teal), NeuN-peaks (salmon), both (dark gray), or neither (light gray)**. Significance symbols denote Bonferroni-adjusted p-values from 2-tailed t-tests for N=18 possible pairwise comparisons, N.S. not significant, * P < 0.05/N, ** P < 0.01/N, *** P < 0.001/N. **(B)** Locus plot candidate SNP with predicted function SNP impact in **cortical excitatory and striatal D1, and D2 MSN** cell types. Genome tracks from top to bottom: **human (h)NeuN+ MACS2 ATAC-seq fold change signal of cortical and striatal brain** regions enriched in Figure 1A. SNP tracks **show lead SNPs aggregated across seven addiction-associated GWAS and the SNPs either in LD with the lead SNPs (Lead SNPs) or independently significant SNPs (LD/ Sig. SNPs). Each SNP is color by increasing red intensity by the degree of LD with a lead SNP. Prioritized candidate causal SNPs by predicted differential cell type activity and overlap with Fullard *et al*. NeuN+ OCRs are plot as (red for Tier A, yellow for Tier B, and teal for Tier C, Methods). Tier A SNP rs7604640 is predicted to have strong ΔSNP effect by CPU-D1 and NAc-D1 CNN models and the bars are colored by the % change in probability active. G**ene annotation tracks **plot** GENCODE **genes from the** GRCh38 **build**. eQTL link tracks of FDR-significant GTEX cis-eQTL from cortical and striatal brain **regions**, and ortholog**s of mouse (m)** putative CREs mapped from **excitatory or striatal neuronal subtypes measured by** cSNAIL ATAC-seq. Cell type colors label cortical excitatory neurons (EXC; red), D1 medium spiny neurons (D1; blue), or D2 medium spiny neurons (D2; green). **(C) Representative importance scores of 50bp flanking either side of the SNP rs7604640 that measure relative contribution of that sequence being active in D1 MSNs. CNN importance score interpretations are shown for effect and non-effect alleles, and the difference in importance scores reveal the relatively more important DNA motif in the effect allele that matches consensus POU1F1 motif overlapping the rs7604640 SNP. The model interprets this POU1F1 motif and a nearby NRF1 motif as contributing to the effect allele having more activity in D1 MSNs.**

One such SNP **from Tier A**, rs7604640, lies in **human** NeuN+ open chromatin specific to striatum 46kb upstream of the *SIX3* locus on chromosome 2. rs7604640 overlaps human orthologs of mouse OCRs in only D1 and D2 neurons and **we predict the effect allele of rs7604640 has an increased probability of open chromatin activity in D1 OCRs of the striatum compared to the non-effect allele (Figure 4B**). rs7604640 is one of many off-lead SNPs identified in the SmokingInitiation GWAS (P_GWAS_ =3.04 x 10^-12^) and is in **high** LD with the SNP rs163522 **(R^2^ = 0.856, P_GWAS_ =1.11 x 10^-11^), which is independently significant from the lead SNP, rs1004787 (R^2^ = 0.630, P_GWAS_ =5.27 x 10^-17^). rs7604640 was reported by HaploRegv4 to overlap a POU1F1 motif** (Ward and Kellis, 2016)**, which our D1 models predict to contribute towards increased probability of being active in D1 MSNs (Figure 4C).** Furthermore, this SNP is a known *cis*-eQTL for the antisense *SIX3-AS1* gene in striatal regions from the Genotype-Tissue Expression (GTEX) project (GTEx Consortium, 2013, 2015; Melé et al., 2015; GTEx Consortium et al., 2017). Anti-sense gene expression is one mechanism of regulating their sense gene (Pelechano and Steinmetz, 2013; Barman et al., 2019), and deletion of the gene *SIX3* has been shown to inhibit development of D2 medium spiny neurons (Xu et al., 2018). Altogether, this evidence formulates the hypothesis that common genetic variant rs7604640 has D1 MSN-specific**, allelic impact on** open chromatin activity in a mouse-human conserved putative CRE regulating the MSN regulator *SIX3*.

**In addition to rs7604640, we report four loci with 1-4 candidate SNPs each in Tier A** that may be putative causal SNPs with cell type-specific activity in addiction-associated traits **(Supplemental Figure 6)**. **The SNPs in these loci all have reported eQTL in frontal cortex** or striatum tissues from GTEx, and they overlap corresponding NeuN+ OCRs and mouse-human orthologous OCRs. In some cases, our prioritized Tier A SNPs were predicted to have ΔSNP effects (Methods) in only striatal MSNs, showcasing our framework’s ability to **predict cell type-specific impact. These SNPs include rs11191352 (P_SmokingInitiation_=2.12 x 10^-7^, Supplemental Figure 6A), rs9826458 (P_RiskyBehavior_= 4.36 x 10^-21^, P_SmokingInitiation_=1.21 x 10^-14^, Supplemental Figure 6B), and rs9844736 (P_RiskyBehavior_= 3.04 x 10^-7^, P_SmokingInitiation_=3.58 x 10^-7^, Supplemental Figure 6C). In a few cases, our models predicted SNPs to have strong** ΔSNP effects across both cortical excitatory and striatal cell types. These include two SNPs in the highly pleiotropic MAPT-CRHR1 locus that are 152bp apart and in perfect LD with each other, rs11575895 and rs62056779 (Supplemental Figure 6D). The prioritized SNPs in the MAPT-CRHR1 locus are genome-wide significant for 5 of the 7 addiction-associated traits (Supplemental Table 1) and the locus has been implicated in other neuropsychiatric traits such as Alzheimer’s Disease (Hoffman et al., 2019; Corces et al., 2020; Ramamurthy et al., 2020). We provide the summary of CNN predictions in these reported loci across all 14,790 analyzed SNPs along with the accompanying annotations that we incorporated into our prioritization of candidate causal SNPs and their predicted cell types (Supplemental Table 1).

## DISCUSSION

In this study, we demonstrate the first analyses integrating **neuronal subtype** OCRs across human and mouse brain epigenomics using **CNN** models to select candidate addiction-associated SNPs acting at putative **neuronal subtype**-specific CREs. We trained **CNN** models to predict **neuronal subtype**-specific activity of OCRs and used the models to predict whether addiction-associated genetic variants in risk loci impact putative CRE function. Our findings link the genetic heritability of addiction-associated behaviors to the OCR profiles of **neuronal subtypes** and brain regions and present specific hypotheses describing how genetic variants may impact gene regulation in addiction-associated behaviors. These analyses in conjunction suggest that genetic variation-associated nicotine, alcohol, and cannabis use behaviors may impact putative CREs in different combinations of excitatory (EXC), D1, and D2 neuronal subtypes. These findings provide a foundation for future investigations into the cell type-specific genetic mechanisms underlying addiction-related traits. More broadly, our cross-species integrative computational framework leverages high-resolution cell-type targeted epigenomics in model organisms to interpret the genetic risk variants of complex traits in humans.

We initially found that addiction-associated genetic variants were enriched in human NeuN+ OCRs of the prefrontal cortex and striatum, known areas involved in addiction and reward circuitry (Volkow et al., 2013; Koob and Volkow, 2016) (**Figure 5A**). Genetic variants associated with SmokingInitiation and Cannabis, initiating behaviors of substance use, were enriched in NeuN+ OCRs of prefrontal areas including DLPFC, VLPFC, and OFC (**Figure 1B**). These OCRs were predicted to be active in cortical excitatory neurons of these brain regions (**Figure 3B**). Addiction-associated genetic variants that enrich in OCRs of cortical excitatory neurons in these areas may reduce corticostriatal activation from prefrontal cortex to inhibit behaviors predisposing the initiation of substance use (Koob and Volkow, 2010, 2016; Volkow et al., 2013; Volkow and Morales, 2015). These genetic variants may contribute to reduced prefrontal self-control reward, leading to behaviors observed in addiction such as impulsivity, reduced satiety, and enhanced motivation to procure drugs (Volkow et al., 2013; Volkow and Morales, 2015). In addition, we found enrichment of striatal NeuN+ OCRs for genetic variants associated with SmokingCessation and DrinksPerWeek (**Figure 1B**). In **Figure 3B**, we showed that these genetic variants are predicted to affect open chromatin in both D1 and D2 MSNs, which are coordinators of mesocorticostriatal dopamine systems (Koob and Volkow, 2010, 2016; Volkow et al., 2013). Genetic variants affecting open chromatin in these MSN subtypes may predispose individuals to increased alcohol use (DrinksPerWeek) or decreased nicotine use (SmokingCessation), perhaps driving the neuroplastic changes in D1 and D2 MSNs observed in human and rodent drug dependence studies (Volkow et al., 1996, 1997, 2003; Wang et al., 1997; Fehr et al., 2008; Cheng et al., 2017; Wilar et al., 2019). While drug addiction has been attributed to various areas of the reward circuit, our investigations into heritable genetic risk for addiction-associated traits unravel how regulatory DNA sequence variation in OCRs of projection neurons in implicated areas link genetic risk to neural circuits to behavior.

**Figure 5.**
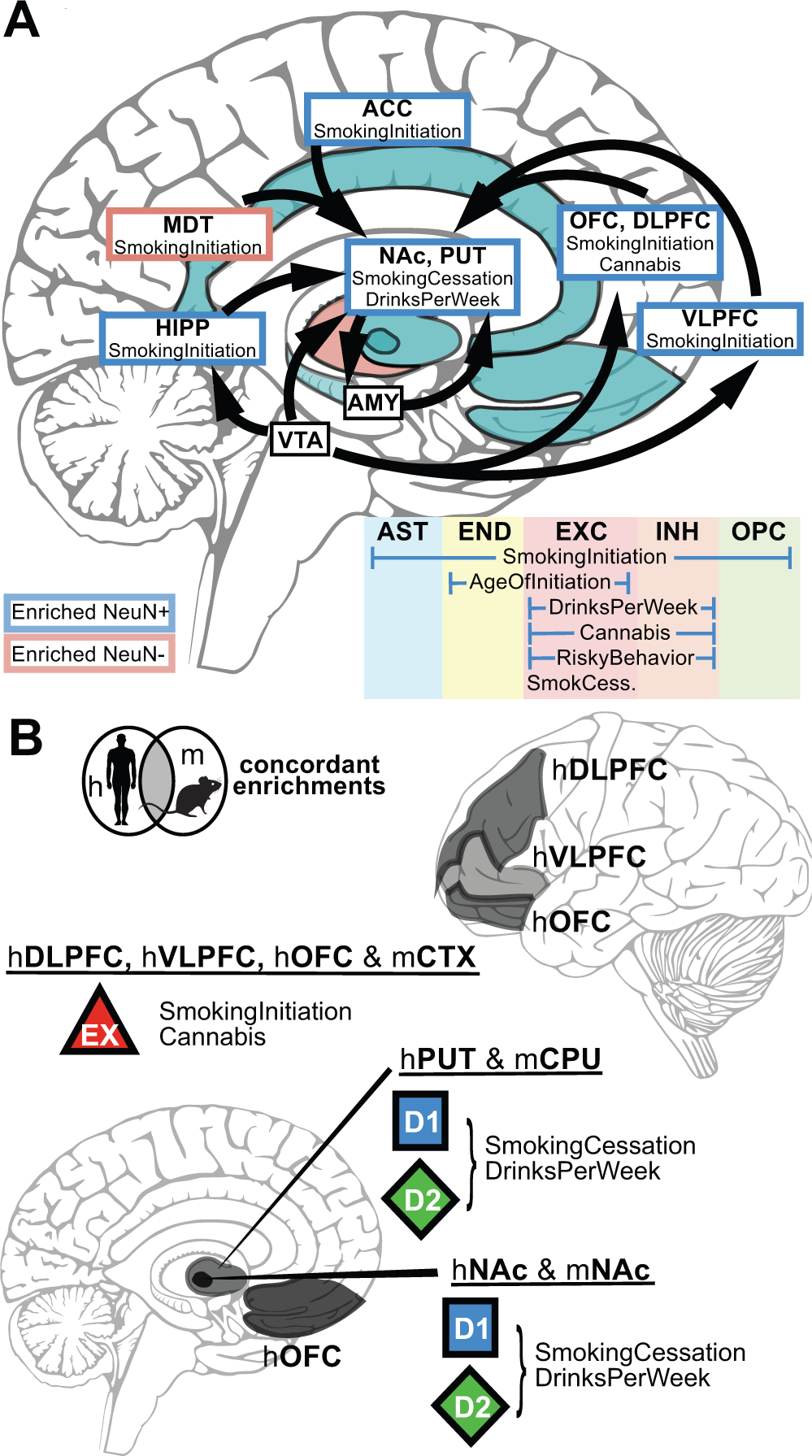
Summary of LDSC GWAS enrichments in human and mouse-human orthologous bulk tissue and cell type open chromatin. **(A)** Schematic of human NeuN-labeled bulk tissue and occipital cortex cell types from **Figure 1** for which addiction-associated genetic variants were significantly enriched (FDR < 0.05) in OCRs. Brain regions are labelled by the cell type that enriched (NeuN+: blue box/shading; NeuN-: red box/shading) spatially along with the trait(s) for which OCRs were found significantly enriched with risk variants. Occipital cortex cell types from **Figure 1C** (same color scheme) are listed along with the trait(s) for which OCRs were found significantly enriched with risk variants. **(B)** Schematic of addiction-associated genetic variants that share enrichments from human brain regions and neuronal subtypes in both human and mouse-human orthologous open chromatin. Brain graphic adapted from Fullard *et al*.(Fullard et al., 2018)

**Since** key component cell types of the reward circuit such as D1 and D2 MSNs have not been profiled for high-quality open chromatin measurements in a human reference genome to our best knowledge, we leveraged high-quality mouse cell type open chromatin measurements using a cross-species OCR mapping framework. We first **conducted ATAC-seq of** MSN **and interneuron subtypes in mouse brain** to identify neuronal subtype-specific OCRs. Then, we used a multiple genome sequence alignment framework to identify the orthologous regions of the human genome. **By leveraging** reference-genome free CRE ortholog mapping tools, we retained high-quality cell type-specific measurements within relevant brain regions of the reward circuit, enabling analysis of cell populations from brain regions where we lack primary human open chromatin profiles. Across these brain regions, we found remarkably concordant enrichments of cell type OCRs between mouse and human profiles as well as shared enrichments between traits (**Figure 5B**). Genetic variants associated with both SmokingInitiation and Cannabis enriched in mouse-human orthologous OCRs of cortical EXC (**Figure 3B**), concordant with enrichments in human cortical NeuN+ OCRs (**Figure 1B**), which were predicted to include EXC neurons (**Figure 4B**). Genetic variants from these two traits showed replicable enrichment in human EXC neuron OCRs of **isocortex and** occipital cortex (**Figure 1C-D**), providing strong evidence that genetic variation in cortical excitatory neuron OCRs confers susceptibility to nicotine and cannabis use behaviors. **The enrichments of genetic variants associated with Cannabis in isocortical IN neuron OCRs (Figure 1C) and mouse-human orthologous OCRs of cortical PV neurons (Figure 3B) suggest that genetic variation in cortical PV neuron OCRs also confer susceptibility of cannabis use behavior.** Within striatal regions, D1 and D2 MSN mouse-human orthologous OCRs enriched for genetic variants of all measured addiction-associated traits (**Figure 2B**), with strongest concordance in human OCRs for genetic variants associated with SmokingCessation and DrinksPerWeek (**Figure 3B, Figure 5B**). The enrichments in conserved OCRs of MSN subtypes in the dorsal striatum and nucleus accumbens unsurprisingly emphasize known roles of MSNs of both areas to drive and maintain addiction behaviors (Ferguson et al., 2011; Ji et al., 2017). **Our validations of enrichments both at the tissue and cell type level across human and human-orthologous OCRs agree with LDSC regression GWAS enrichments of non-coding regions around differentially expressed genes in DLPFC** and NAc measured from postmortem human subjects who were diagnosed with opioid use disorder vs. neuropsychiatric controls **(Seney et al., 2020)**. Due to the conservation of reward circuit between mouse and human, our approach is able to unravel the cell types in which genetic variation at the epigenome level predisposes addiction-related traits even from measurements in organisms that have not been exposed to addictive substances. Further, this level of OCR conservation is present at the level of excitatory cell types in cortical brain regions (cite). This may explain why we found enriched cell types in occipital cortex (Figure 1D), which is not well-defined for its role in addiction-related traits.

In an orthogonal approach to mapping mouse-human orthologous OCRs, **we devised and trained convolutional neural network (CNN) models to classify the neuronal subtype membership of mouse and human NeuN+ OCRs in order to refine GWAS enrichments of bulk tissue to the major neuronal subtypes of cortex and striatum. This approach can provide further validation for enrichments of human and mouse-human orthologous OCRs in cell types and tissues.** Refinement of NeuN+ OCRs revealed that addiction-associated traits enriched for two clusters of cell types and brain regions. The first group, which displays concordant cortical excitatory enrichments between human and mouse, consists of SmokingInitiation and Cannabis (**Figure 3B**), and the second group, which displays concordant D1 and D2 MSN enrichments, consists of SmokingCessation and DrinksPerWeek. A draw-back of assigning human NeuN+ OCR membership to individual cell types lies in the considerably low representation of interneurons in both cortical and striatal neuron populations - as low as 12% in neocortex (Beaulieu, 1993; Lefort et al., 2009) and ∼5% in striatum (Tepper and Koós, 2017; Krienen et al., 2019). NeuN+ open chromatin profiles alone do not always capture OCRs unique to rare interneurons, some of which had OCRs identified by human single-cell assays and mouse-human orthologs enriched for addiction GWAS variants (**Figure 1C, Figure 2B**). As a result, we did not train **CNN** models for PV, SST, or VIP interneurons. However, the striking enrichments of OCRs from certain interneuron populations for addiction GWAS variants begin to demonstrate these populations’ roles in the addiction neural circuits (Bracci et al., 2002; Lansink et al., 2010; Wiltschko et al., 2010; Ribeiro et al., 2018; Jiang et al., 2019; Lee et al., 2020; Schall et al., 2020).

The overall concordance of enrichments across human and mouse-human orthologous OCRs suggests a conserved regulatory **code** between mouse and human CREs. Correspondence in the neural circuitry has been well-appreciated between human studies and rodent models of addiction (Berke and Hyman, 2000; Koob and Volkow, 2016; Farrell et al., 2018), and our study further demonstrates that mouse-human conserved OCRs may explain considerable heritability of addiction-associated traits. This makes animal models suitable not only for studying the neural circuits of addiction but also cell-type-specific gene-regulatory mechanisms of addiction.

We used several selection criteria along with **CNN** models to predict the functional impact of genetic variants **associated with** addiction-**related** traits (**Figure 5, Supplemental Figure 5, Supplemental Table 1)**. The fine-mapping pipeline described effectively narrows down a set of 14,790 SNPs to a putatively functional set of **55 Tier A candidate causal SNPs** that can be experimentally tested to determine which brain regions and neuronal subtypes they would have function in. The candidate functional SNPs that our models prioritize demonstrate how a candidate SNP within a locus, such as rs7604640 (**Figure 4B**), might act in **distinct** neuronal subtypes and brain regions. **Cell type-and brain region-specificity adds complexity to identifying how genetic variation may alter gene regulation to predispose individual to addiction-associated traits. Our approach often reported one to four candidates per loci, even in stretches of SNPs in perfect LD such as the MAPT-CRHR1 locus (Supplemental Figure 5D)**. This reflects the idea that many SNPs in the same loci are significantly associated with addiction due to LD with **only one or a few** causal SNPs and **are unlikely to** influence addiction-associated genetic predisposition. **We report many** candidate SNPs that **also** overlap mouse-human orthologs from the same predicted cell type raise the idea that altering the conserved regulatory DNA sequence may be a mechanism of cell type-specific gene regulatory tuning in a population or even across species (Gjoneska et al., 2015).

Our study depends solely on assays of open chromatin as a proxy for putative CREs. Epigenetic assays for chromatin conformation, histone modifications, and methylation would further inform how putative CREs regulate nearby gene expression. While eQTL studies do not control for inflated associations due to LD and report gene expression differences from bulk tissue, we do note that our approach prioritizes several SNPs known to be *cis-*eQTLs in relevant brain regions, which indirectly affirms our framework’s ability to select SNPs with functional impacts on gene regulation. **Although *cis*-EQTLs are often not cell type-or tissue-specific, our findings of risk loci in brain regions implicated in addiction-related traits reflect a strength of our approach in discerning brain-specific signal.** In order to validate our predictions, it will be necessary to further investigate candidate genetic variants such as rs7604640 (**Figure 4B**) in future studies using **a fluorescence reporter assay or *in situ* hybridization studies. These methods can measure regulatory activity differences between risk and non-risk alleles to verify our predictions of SNP impact on putative CREs and indicate whether the reported differences in regulatory activity are cell type-specific.** The candidate SNPs we identified provide possible mechanisms linking differences in genetic make-up with the genes, cell types, and brain regions that could influence addiction and substance use behaviors (**Figure 4**).

## MATERIALS & METHODS

### ATAC-seq data processing pipeline

We processed raw FASTQ files of ATAC-seq experiments with the official ENCODE ATAC-seq pipeline (Landt et al., 2012) accessed by https://github.com/ENCODE-DCC/atac-seq-pipeline. We ran this pipeline using the mm10 genome assembly for mouse and the hg38 genome for human with the following settings: smooth_win = 150, multimapping = 0, idr_thresh = 0.1, cap_num_peak = 300,000, keep_irregular_chr_in_bfilt_peak = true. We grouped biological replicates when processing data to obtain individual de-duplicated, filtered bam files and reproducible (IDR) peaks for each condition. Unless otherwise stated, we used the optimal reproducible set of peaks for downstream analyses. We removed samples that had low periodicity indicated by ENCODE quality control metrics and reprocessed the remaining replicates with the pipeline.

### Publicly available datasets

Fullard *et al*. NeuN-sorted ATAC-seq of human postmortem brain (Fullard et al., 2018): We identified OCRs overlapping addiction-related variants through analysis of human postmortem brain ATAC-seq in which cells were sorted into NeuN-positive and NeuN-negative groups via fluorescence activated nuclei sorting (FANS); the brain regions we used were dorsolateral prefrontal cortex (DLPFC), orbitofrontal cortex (OFC), ventrolateral prefrontal cortex (VLPFC), anterior cingulate cortex (ACC), superior temporal gyrus (STC), inferior temporal gyrus (ITC), primary motor cortex (PMC), insula (INS), primary visual cortex (PVC), amygdala (AMY), hippocampus (HIP), mediodorsal thalamus (MDT), nucleus accumbens (NAc), and putamen (PUT). We downloaded data from the Sequence Read Archive (SRA) through Gene Expression Omnibus (GEO) accession #GSE96949. We separated samples by cell type and reprocessed them with the ENCODE pipeline as detailed above, aligning reads to hg38. We used the “optimal reproducible peaks” for each cell type and brain region as foregrounds in GWAS LDSC enrichment with the Honeybadger2 OCR set as the background set (see LDSC Regression GWAS Enrichment Backgrounds).

Corces *et al*. human isocortex, striatum, hippocampus, and substantia nigra single cell chromatin accessibility profiling **(Corces et al., 2020)**: We downloaded 24 clusters of IDR peaks in BED format through GEO accession #GSE147672. These clusters represent cell populations defined by Corces *et al* from the measured brain regions. We assigned clusters to cell populations as described in Corces *et al*: astrocyte (clusters 13, 17), hippocampal excitatory (clusters 3-4), isocortical astrocyte (cluster 15), isocortical excitatory (cluster 1), isocortical inhibitory (cluster 11), microglia (cluster 24), neuron (cluster 7), nigral astrocyte (cluster 14), nigral neurons (clusters 5-6), nigral oligodendrocyte precursor (cluster 10), oligodendrocyte (clusters 19-23), oligodendrocyte precursor (clusters 8-9), striatal astrocyte (cluster 16), and striatal inhibitory cells (clusters 2, 12). We did not include cluster 18, which corresponds to a doublet. We merged coordinates from clusters assigned to the same cell types to define foreground sets for LDSC regression GWAS enrichment. We merged the foreground sets with the Honeybadger2 OCR set to define the background set (LDSC regression GWAS Enrichment Backgrounds).

Lake *et al*. human occipital cortex scTHS-seq (Lake et al., 2018): We downloaded BED-formatted cell type-specific differential OCRs from occipital cortex scTHS-seq of excitatory neurons (EXC), inhibitory neurons (IN), astrocytes (AST), endothelial cells (END), oligodendrocyte precursor cells (OPC), oligodendrocytes (OLI), and microglia (MIC) from the GEO subseries #GSE97887. We used the hg38 OCR coordinates as foregrounds in LDSC regression GWAS enrichment with the Honeybadger2 OCR set as the background set (LDSC regression GWAS Enrichment Backgrounds).

Mo *et al*. mouse INTACT-sorted nuclei ATAC-seq (Mo et al., 2015): We downloaded FASTQ files of *R26-CAG-LSL-Sun1-sfGFP-Myc* transgenic mouse lines for cell type-specific ATAC-seq performed using the INTACT method from the accession #GSE63137. *Mo et al.* isolated INTACT-enriched nuclei from three cell types: excitatory neurons (EXC, *Camk2a-cre*), vasoactive intestinal peptide neurons (VIP, *Vip-cre*), and parvalbumin neurons (PV, *Pvalb-cre*). We reprocessed the data with the Kundaje Lab open chromatin pipeline using the mm10 genome (https://github.com/kundajelab/atac_dnase_pipelines). We mapped reproducible mouse ATAC-seq peaks for each cell type to hg38 using halLiftover with the 12-mammals Cactus alignment (Paten et al., 2011; Hickey et al., 2013) followed by HALPER (Zhang et al., 2020) (Mapping mouse OCR orthologs) to produce a foreground set of orthologous human sequences for LDSC regression GWAS enrichment (Finucane et al., 2018). We mapped the ENCODE mm10 DNaseI-hypersensitive peak set (Yue et al., 2014) to hg38 (Mapping mouse OCR orthologs) and used successfully mapped hg38 orthologs of mm10 OCRs a background set for mouse foreground enrichments. Furthermore, we used this dataset to evaluate differential accessibility in cSNAIL-INTACT PV and PV-negative ATAC-seq samples and develop **convolutional neural network** models of cell type-specific open chromatin (see Methods below).

Human negative control foregrounds (ENCODE Project Consortium, 2012; Thurman et al., 2012; Davis et al., 2018; Cannon et al., 2019): We downloaded raw ATAC-seq profiles of human adult female and male stomach ATAC-seq generated by Snyder *et al*. (ENCSR337UIU, ENCSR851SBY, respectively), female human embryonic liver DNase-seq generated by Stamatoyannopoulos *et al*. (ENCSR562FNN), and human embryonic lung DNase-seq generated by Stamatoyannopoulos *et al*. (ENCSR582IPV) from https://www.encodeproject.org/. We processed these files using the ENCODE pipeline as detailed above to obtain optimal reproducible hg38 peaks. We also downloaded BED files of human adipocyte and preadipocyte ATAC-seq profiles generated by Cannon *et al*. from GEO accession number #GSE110734. We mapped these BED coordinates from hg19 to hg38 using liftOver to define negative control foregrounds for human LDSC regression GWAS enrichment. We merged the human negative control foregrounds and Fullard *et al*. foregrounds with the Honeybadger2 OCR set to define the background for human negative control foreground enrichments.

Human-orthologous negative control foregrounds (Liu et al., 2019a): We also downloaded raw ATAC-seq data profiled in female mouse kidney, female mouse liver, and male mouse lung generated by Liu *et al*. from SRA accession #SRP167062 to define human-orthologous negative control foregrounds. We processed these files using the ENCODE pipeline as detailed above to get optimal reproducible peaks. We mapped optimal reproducible peaks from mm10 to hg38 using halLiftover with the 12-mammals Cactus alignment followed by HALPER (Mapping mouse OCR orthologs) to define negative control foregrounds for human-orthologous LDSC GWAS enrichments. We merged all human orthologous foregrounds with the human orthologs of the ENCODE mm10 DNaseI-hypersensitive peak set to define a background for human-orthologous LDSC GWAS enrichments.

### Mapping mouse open chromatin region (OCR) orthologs

We employed halLiftover (Hickey et al., 2013) with the 12-mammals Cactus alignment (Paten et al., 2011) followed by HALPER (https://github.com/pfenninglab/halLiftover-postprocessing) (Zhang et al., 2020) to map mm10 mouse reproducible OCRs to hg38 human orthologs in order to perform LDSC regression GWAS enrichment. The Cactus multiple sequence alignment file (Paten et al., 2011) has 12 genomes, including mm10 and hg38, aligned in a reference-free manner, allowing us to leverage multi-species alignments to confidently identify orthologous regions across species. halLiftover uses a Cactus-format multiple species alignment to map BED coordinates from a query species to orthologous coordinates of a target species, and HALPER constructs contiguous orthologs from the outputs of halLiftover (Zhang et al., 2020). We ran the orthologFind.py function from HALPER on the outputs of halLiftover using the following parameters: -max_frac 5.0 -min_frac 0.05 -protect_dist 5 -narrowPeak -mult_keepone. In general, 70% of mouse brain ATAC-seq reproducible peaks were able to be mapped to confident human orthologs. To map the ENCODE mm10 mouse DHS background, which does not contain summit information, to hg38 we used the mouse coordinates of position with the most species aligned in a region to define the summit. Only for the mm10 mouse DHS background set, for which a significant proportion of regions could not be confidently mapped to hg38, we flanked the original assembly coordinates by 300 bp to increase OCR mapping from 54% to 64%.

### LDSC Regression GWAS Enrichment Backgrounds

We found that LDSC regression GWAS enrichment analysis is sensitive to the selected background set of matched regions. To construct appropriate background sets for each GWAS enrichment, we used the ENCODE and RoadMap Honeybadger2(Roadmap Epigenomics Consortium et al., 2015) and Mouse DHS peak sets for the respective human and mouse-based open chromatin GWAS enrichment. The Honeybadger2 set, downloaded from https://personal.broadinstitute.org/meuleman/reg2map/, consists of DNaseI-hypersensitive OCRs across 53 epigenomes consisting of promoter, enhancer, and dyadic regions. Honeybadger2 is an appropriate epigenetic reference for enrichment of cell type-specific open chromatin from various foregrounds such as the Fullard *et al*. and Lake *et al*. Honeybadger2 regions allow the LDSC algorithm to properly account for the heritability from OCRs of a particular cell type or regions rather than because they tend to be more conserved, are enriched for ubiquitously active transcription factor motifs, or other factors distinguishing open chromatin from heterochromatin. The human orthologs of the ENCODE Mouse DHS peak set, downloaded through the ENCODE ATAC-seq pipeline at http://mitra.stanford.edu/kundaje/genome_data/mm10/ataqc/mm10_univ_dhs_ucsc.bed.gz, is a set of peaks combined from mouse DNaseI-hypersensitivity

OCRs from ENCODE and provides a background for human orthologs of mouse OCRs. The mm10 mouse DHS regions were mapped to hg38 as described in **Mapping mouse OCR orthologs**. For each respective foreground-background pairing, the foreground regions were merged with the background reference to ensure the background always contained the foreground set. The mouse background was used to calculate the significance of the relationship between mouse OCRs and GWAS for addiction-associated traits to control for a possible association between the degree to which a region is conserved and its likelihood in influencing the predisposition to an addiction-associated trait.

### GWAS enrichment with partitioned LD score regression analysis

We computed the partitioned heritability of CREs for GWAS variants using the LDSC regression pipeline for cell type-specific enrichment as outlined in https://github.com/bulik/ldsc/wiki/Cell-type-specific-analyses (Bulik-Sullivan et al., 2015b). We downloaded the GWAS summary statistics files and processed them with the LDSC munge_sumstats function to filter rare or poorly imputed SNPs with default parameters. We munged the summary statistics files for HapMap3 SNPs excluding the MHC regions downloaded at http://ldsc.broadinstitute.org/static/media/w_hm3.noMHC.snplist. zip. We inspected GWAS file to ensure the effect allele, non-effect allele, sample size, p-value, and signed summary statistic for each SNP in each GWAS were included and appropriate for LDSC. The addiction-associated GWAS measure genetic predisposition for age of smoking initiation (AgeofInitiation) (Liu et al., 2019b), heaviness of smoking (CigarettesPerDay) (Liu et al., 2019b), having ever regularly smoked (SmokingInitiation) (Liu et al., 2019b), current versus former smokers (SmokingCessation) (Liu et al., 2019b), alcoholic drinks per week (DrinksPerWeek) (Liu et al., 2019b), cannabis consumption (Cannabis) (Pasman et al., 2018), and risk tolerance (RiskyBehavior) (Karlsson Linnér et al., 2019). GWAS traits related to addiction include multisite chronic pain (ChronicPain) (Johnston et al., 2019) and number of coffee cups drank per data (CoffeePerDay) (Coffee and Caffeine Genetics Consortium et al., 2015). Other addiction-related traits come from underpowered GWAS including opioid dependence (OpioidDep) (Cheng et al., 2018), cocaine dependence (CocaineDep) (Cabana-Domínguez et al., 2019), and diagnosis of obsessive-compulsive disorder (OCD) (International Obsessive Compulsive Disorder Foundation Genetics Collaborative (IOCDF-GC) and OCD Collaborative Genetics Association Studies (OCGAS), 2018). GWAS from strong brain-related traits used are schizophrenia risk (Schizophrenia)(Schizophrenia Working Group of the Psychiatric Genomics Consortium, 2014), highest level of educational attainment (EduAttain) (Lee et al., 2018), and sleep duration (SleepDuration) (Dashti et al., 2019). The non-brain related traits measure genetic liability for lean body mass (LBM) (Zillikens et al., 2017), bone mineral density (BMD) (Kemp et al., 2017), and coronary artery disease (CAD) (Howson et al., 2017).

We estimated LD scores for each foreground set and corresponding background set with the LDSC regression pipeline make_annot and ldsc functions using hg38 1000 Genomes European Phase 3 European super-population (1000G EUR) cohort plink files downloaded from https://data.broadinstitute.org/alkesgroup/LDSCORE/GRCh38/. An example of an ATAC-seq optimal set of reproducible peaks mapped to hg38 in narrowPeak format is annotated with 1000G EUR LD scores using the following call:

python make_annot.py \

--bed-file optimal_peak.narrowPeak.gz \

--bimfile 1000G.EUR.hg38.${chr}.bim \

--annot-file foreground.${chr}.annot

We downloaded the baseline v1.2 files for cell type-specific enrichment in hg38 coordinates from the same link above as well as the corresponding weights ‘weights.hm3_noMHC’ file excluding the MHC region from https://data.broadinstitute.org/alkesgroup/LDSCORE/. HapMap SNPs and corresponding weights file used in the LDSC analyses only refer to the SNP rsIDs, rather than genomic coordinates, so only the baseline and LD statistics used to annotate the foreground and background files must be in hg38 coordinates. In accordance with the LDSC regression script input format, we created an ‘enrichment.ldcts’ file listing the annotated foreground/background pair for each foreground set. We estimated the partitioned heritability using the ldsc function, which integrates the foreground and background LD score estimates, munged GWAS SNP data, baseline variant data, and variants weights. The final function call to GWAS enrichment is as follows:

python ldsc.py --h2-cts $Munged_GWAS \

--ref-ld-chr baseline_v1.2/baseline. \

--w-ld-chr weights.hm3_noMHC. \

--ref-ld-chr-cts enrichment.ldcts \

--out $Output_Label

The pipeline produced LD score regression coefficient, coefficient error, and coefficient p-value estimates. We adjusted for multiple testing using the false discovery rate on p-values of the LD score regression coefficients (alpha = 0.05) on all 18 GWAS traits intersected on within the same foreground/background set. A significant FDR-value indicates enrichment of the foreground genomic regions for GWAS SNPs relative to the background. Lastly, we computed genetic correlations in **Supplemental Figure 1A** between GWAS of addiction-associated traits using the pre-munged summary statistics as described by Bulik-Sullivan *et al*. (Bulik-Sullivan et al., 2015a)

### Bulk tissue ATAC-seq

To augment and compare to mouse cell type-specific ATAC-seq datasets generated in this study, we also performed bulk tissue ATAC-seq from cortex (CTX) and dorsal striatum/nucleus accumbens (CPU) of 7- and 12-week-old C57Bl/6J mice (N = 2 each age) as described in Buenrostro *et al*., 2015(Buenrostro et al., 2015) with the following minor differences in buffers and reagents. We euthanized mice with isoflurane, rapidly decapitated to extract the brain, and sectioned it in ice-cold oxygenated aCSF (119mM NaCl, 2.5 mM KCl, 1mM NaH_2_PO_4_(monobasic), 26.2mM NaHCO_3_, 11mM glucose) at 200-micron sections on a vibratome (Leica VT1200). We further micro-dissected sections for cortex and dorsal striatum on a stereo microscope and transferred dissected regions into chilled lysis buffer (Buenrostro et al., 2015). We dounce homogenized the dissected brains in 5mL of lysis buffer with the loose pestle (pestle A) in a 15mL glass dounce homogenizer (Pyrex #7722-15). We washed nuclei lysate off the pestle with 5mL of lysis buffer and filtered the nuclei through a 70-micron cell strainer into a 50mL conical tube. We washed the dounce homogenizer again with 10mL of BL buffer and transferred the lysate through the 70-micron filter (Foxx 1170C02). We pelleted the 20 mL of nuclei lysate at 2,000 x *g* for 10 minutes in a refrigerated centrifuge at 4°C. We discarded the supernatant and resuspended the nuclei in 100-300 microliters of water to approximate a concentration of 1-2 million nuclei/ mL. We filtered the nuclei suspension through a 40-micron cell strainer. We stained a sample of nuclei with DAPI (Invitrogen #D1206) and counted the sample to measure 50k nuclei per ATAC-seq transposition reaction. The remaining steps follow the Buenrostro *et al*., 2015 (Buenrostro et al., 2015) protocol for tagmentation and library amplification. We shallowly sequenced barcoded ATAC-seq libraries at 1-5 million reads per sample on an Illumina MiSeq and processed individual samples through the ENCODE pipeline for initial quality control. We used these QC measures (clear periodicity, library complexity, and minimal bottlenecking) to filter out low-quality samples and re-pooled a balanced library for paired-end deep sequencing on an Illumina NextSeq to target 30 million uniquely mapped fragments per sample after mitochondrial DNA and PCR duplicate removal. These raw sequencing files entered processing through the ENCODE ATAC-seq pipeline as above by merging technical replicates and grouping biological replicates by brain region for each pipeline run.

### Cre-Specific Nuclear-Anchored Independent Labeling (cSNAIL) virus procedures

The cSNAIL genome (pAAV-Ef1a-DIO-Sun1-Gfp-WPRE-pA) contains *loxP* sites to invert the *Sun1-Gfp* fusion gene and integrate into the nuclear membrane of cells expressing the *Cre* gene, allowing these cell populations to be profiled for various genomic assays (Lawler et al, 2020 in press J. Neuro). We packaged the cSNAIL genome with AAV variant PHP.eB (pUCmini-iCAP-PHP.eB) in AAVpro(R) 293T cells (Takara, cat #632273). Viviana Gradinaru provided us with the pUCmini-iCAP-PHP.eB (http://n2t.net/addgene:103005; RRID: Addgene 103005) (Chan et al., 2017). We precipitated viral particles with polyethylene glycol, isolated with ultracentrifugation on an iodixanol density gradient, and purified in PBS with centrifugation washes and 0.2µM syringe filtration. We injected each mouse with 4.0 x 10^11^vg into the retro-orbital cavity under isoflurane anesthesia. We allowed the virus to incubate in the animal for 3-4 weeks to reach peak expression. We closely monitored the health of the animals throughout the length of the virus incubation and did not note any concerns.

### cSNAIL nuclei isolation

On the day of the ATAC-seq experiments, we dissected brain regions from fresh tissue and extracted nuclei in the same manner as described for bulk tissue experiments. Then, we sorted the nuclei suspension into Sun1GFP+ (Cre+) and Sun1GFP-(Cre-) fractions using affinity purification with Protein G Dynabeads (Thermo Fisher, cat. 10004D). A pre-clearing incubation with beads and nuclei for 10-15 minutes removes effects from non-specific binding events. Next, we incubated the remaining free nuclei with anti-GFP antibody (Invitrogen, #G10362) for 30 minutes to bind Sun1GFP. Finally, we added new beads to the solution to conjugate with the antibody and incubated the reaction for an additional 20 minutes. The pre-clear step and all incubations took place in wash buffer (0.25M Sucrose, 25mM KCl, 5mM MgCl_2_, 20mM Tricine with KOH to pH 7.8, and 0.4% IGEPAL) at 4°C with end-to-end rotation. After the binding process, we separated bead-bound nuclei on a magnet, washed three times with wash buffer, and filtered through a 20µM filter to ensure purity. We resuspended nuclei in nuclease-free water for input into the ATAC-seq tagmentation reaction. We performed nuclei quantification and tagmentation in the same manner described for bulk tissue ATAC-seq above. We list in the table below the number of animals, the genotypes, and which regions collected for ATAC-seq experiments in this study. N=2 *Pvalb-cre* samples from CPU/NAc region had received a sham surgery with saline injection into the external globus pallidus 5 days before they were sacrificed. **The background for all transgenic mice is C57BL/6J. SST-Cre mice were homozygous for the transgene while PValb-2a-Cre, D1-Cre, and A2a-Cre mice were heterozygous for the transgene (Lawler et al., 2020)**. N=2 *Drd1-cre* samples from both CPU and NAc regions had received headcap surgeries 3 weeks before they were sacrificed. Both *Pvalb-cre* and *Drd1-cre* were overall healthy at time of sacrifice.

| Genotype | Replicates | Sex<br>(Female /Male) | Region and Replicate<br>per region | Cell<br>type |
| --- | --- | --- | --- | --- |
| C57BL/6 WT | N=4 | 2 F, 2 M | CTX= 4, CPU/NAc = 4 | bulk |
| *Pvalb-cre* | N=5 | 3 F (CTX)<br>1 F, 1 M (CPU/NAc) | CTX= 3, CPU/NAc = 2 | PV |
| *Sst-cre* | N=2 | 1 F, 1 M | CTX= 2, CPU/NAc = 2 | SST |
| *Drd1-cre* | N=2 | 2 F | CPU=2, NAc=2 | D1 |
| *Adora2a-cre* | N=2 | 2 F | CPU=2, NAc=2 | D2 |

### cSNAIL Cell Type Specificity

We created a consensus set of non-overlapping IDR peaks from the ATAC-seq pipeline for cSNAIL ATAC-seq and Mo *et al*. INTACT samples (Tissue: Ctx, CPU, and NAc ; Celltype: EXC, PV, SST, VIP, D1, D2). We extended the peak set 200bp up- and down-stream, count overlapping fragments with Rsubread v2.0.1 using the de-duplicated BAM files from the pipeline(Liao et al., 2014), and created with DESeq2 v1.26.0 a variance-stabilized count matrix aware of experimental Group (combination of Tissue and Celltype) with ∼Group (Love et al., 2014). We plotted the principle component analysis in **Supplemental Figure 3A** for the first two components with this variance-stabilized count matrix. We used Deeptools v3.5.0 to convert the same BAM files to normalized bigWig files and average over replicates of the same Group (Ramírez et al., 2016). We plotted the tracks using pyGenomeTracks v3.5 around marker genes for each cell type (*Slc17a7*, *Drd1*, *Adora2a*, *Pvalb*, *Sst*, *Vip*) (Ramírez et al., 2018) **Supplemental Figure 3B**. We computed the mean accessibility for each Group 2kb up- and down-stream the transcription start sites (TSS) and correlated log_10_ (TSS accessibility + 1) with gene expression log_10_(meta gene counts + 1) of Drop-Seq annotated cell types from prefrontal cortex and striatum(Saunders et al., 2018). We used the Saunders *et al*. tissue subcluster metagene profiles (sum of gene expression in all cells) and summed subclusters to cluster-level metagene profiles. Most tissue cluster metagene profiles corresponded to cSNAIL ATAC-seq celltype and tissue profiles, with the exception of cSNAIL cortical PV+ samples were matched to Saunders *et al*. cortical MGE+ interneuron clusters.

### Convolutional Neural Network models for CRE cell type classification

We trained a set of convolutional neural network (CNN) models to learn the regulatory **code** of a given cell type from the DNA sequences underlying the cell type’s OCRs. The models take in one-hot encoded 501bp genomic sequences **to predict 1 for an OCR or 0 for non-OCR sequence**. Positive sequences were centered on IDR peak summits that are annotated to be in introns and distal intergenic regions and negative sequences are approximately ten times the number of positives sequences that are G/C-matched and not overlapping IDR peaks. We excluded promoters (defined as within 5,000bp from the TSS) and exons because distal sequences have been shown to confer more cell type-specificity and be more predictive of expression levels of regulated genes (Roadmap Epigenomics Consortium et al., 2015). We constructed the negative set by first building a sequence repository $BGDIR according to https://bitbucket.org/CBGR/biasaway_background_construction/src/master/ from the mouse mm10 genome using 501bp sequences. Then we used the biasaway (Worsley Hunt et al., 2014; Khan et al., 2020) command-line interface to generate negative sequences with the matching nucleotide distribution along a sliding window of the 501bp IDR peak sequence: biasaway c --foreground $FGFASTA --nfold 10 --deviation 2.6 -- step 50 --seed 1 –winlen 100 --bgdirectory $BGDIR

We employed a 5-fold cross validation chromosome hold-out scheme to train 5 models per set of IDR peaks to ensure stable and consistently learned regulatory patterns. A model that was training a fold did not see sequences during training from the validation set for that fold, and no model saw the test set until final model performance evaluation. Sequences from these chromosomes were used as the validation set for each fold: fold1: {chr6, chr13, chr21} fold2: {chr7, chr14, chr18} fold3: {chr11, chr17, chrX} fold4: {chr9, chr12} fold5: {chr10, chr8}.

We used sequences from chromosomes {chr1, ch2, chr19} for the test set. We trained the models with Keras v2.3.0-tf (https://keras.io/) implemented through Tensorflow v2.2.0 and used stochastic gradient descent (SGD) with Nesterov **momentum to minimize the binary cross entropy loss and learn model parameters. All models used the same CNN** architecture after a grid-search of hyperparameters found stable and high validation performance by area under the precision-recall curve (auPRC) in an architecture with five Conv1D layers (kernel_size = 11, filters = 200, activation= ‘relu’, kernel_regularizer=l2(1e-10)) sandwiched between four Dropout layers (rate = 0.25), then one MaxPooling1D layer (pool_size = 26, strides = 26), one Flatten layer, one Dense layer (units = 300, activation=‘relù, kernel_regularizer=l2(1e-10)), one Dropout layer **(rate = 0.25), a final output Dense layer (units=1, activation = ‘sigmoid’, kernel_regularizer=l2(1e-10)), and a final Dropout layer (rate = 0.25) before the sigmoid output layer**. We applied the One-Cycle-Policy (OCP) with linear cyclical learning rate and momentum between a base and max rates as described previously (Smith, 2018) to train each fold with batch_size= **1000**, epochs = 23, num_cycles = 2.35, base_learning_rate = 1e-2, max_learning_rate = 1e-1, base_momentum = .85, max_momentum = 0.99. **With these hyperparameters, we** trained models across folds to predict positive OCRs of all **measured** cell types against an approximately 1:10 positive:negative class **ratio**. We computed classifier performance metrics including weighted accuracy (using threshold = 0.5), weighted f1_score (using threshold = 0.5), area under receiver operating characteristic (auROC), and area under precision-recall curve (auPRC). **Given the class imbalance, we selected the reported hyperparameters that maximize the validation auPRC at a threshold of 0.5. We report the test performance auPRC, F1 score, and false positive rate on 10X nucleotide-content matched negatives in Supplemental Figure 5A. We** provide both the scripts and trained Keras models at https://github.com/pfenninglab/addiction_gwas_enrichment.

### Interpretation of Convolutional Neural Network models

**To ensure that the classification task decisions relied on biological sequence signatures and not artifacts, we performed model interpretation using Deep SHAP v0.37.0** (Štrumbelj and Kononenko, 2014; Shrikumar et al., 2017) **and TF-MoDISco** (Shrikumar et al., 2018)**. For a random subsample of 2,000 true positive sequences from the validation set, we generated per base importance scores and hypothetical importance scores relative to a reference set of 500 true negative sequences from the validation set. These scores describe the contribution of each base toward a positive model classification, which is a predicted OCR in the given cell type. TF-MoDISco is an importance score-aware method that clusters commonly important subsequences, called “seqlets”, to define the learned motifs of the model. We ran TF-MoDISco v0.4.2.3 with the options sliding_window_size=11, flank_size=3, min_seqlets_per_task=3000, trim_to_window_size=11, initial_flank_to_add=3, final_flank_to_add=3, kmer_len=7, num_gaps=1, and num_mismatches=1. The resulting motifs were filtered to remove rare patterns with fewer than 100 supporting seqlets. Then, the motifs were visualized and associated with known motifs using Tomtom** (Gupta et al., 2007) **version 5.3.3 with the HOCOMOCO v11 FULL database and default parameters (Supplementary Table 2)**.

### Machine learning cell type-specific prioritization of *Fullard et al.* NeuN+ ATAC-seq peaks

We used CNN model scores to classify whether a peak from *Fullard et al.* NeuN+ open chromatin data is active in a neuronal subtype {EXC, D1, D2}. We took NeuN+ IDR “optimal peaks” from regions significantly enriched for addiction-associated traits {OFC, VLPFC, DLPFC, ACC, STC, PUT, NAc, **Figure 1A**}, extracted 501bp DNA sequences of each centered on the summit, and scored each peak with cell type-specific machine learning models trained with the appropriate tissue context (e.g., score cortical NeuN+ peaks with a model trained with cortical EXC cell type). We averaged scores across model folds from the same cell types and classified NeuN+ peaks with scores greater than 0.5 as active in that cell type**, as this threshold was the most discriminative in classifying positive validation set sequences (Supplemental Figure 5B)**. We defined these CNN-prioritized peaks as foregrounds for LDSC regression GWAS enrichment analyses as described above. We created a consensus set of peaks merging all model-prioritized peaks and the Honeybadger2 set of OCRs to be the matched background, and we performed GWAS enrichment and computed FDR on all 18 GWAS traits (only enrichments for addiction-associated GWAS shown, **Figure 3**).

### Addiction-associated GWAS processing and cell type-specific candidate selection

We collected the addiction-associated SNPs by submitting the summary statistics files for the seven addiction-associated traits {AgeofInitiation (Liu et al., 2019b), CigarettesPerDay (Liu et al., 2019b), SmokingInitiation (Liu et al., 2019b), SmokingCessation (Liu et al., 2019b), DrinksPerWeek (Liu et al., 2019b), Cannabis (Pasman et al., 2018), RiskyBehavior (Karlsson Linnér et al., 2019)} to the FUMA webserver (Watanabe et al., 2017). FUMA computed LD R^2^ based on the 1000 Genomes European (1000G EUR) super-population reference (1000 Genomes Project Consortium et al., 2015) via PLINK (Purcell et al., 2007), linked GWAS-significant lead SNPs to off-lead SNPs in LD with the lead, and annotated functional consequences of genetic variants via ANNOVAR based on ENSEMBL build 85 human gene annotations (Wang et al., 2010) (**Figure 1A**). ANNOVAR functional gene annotations for a SNP are as defined in the primary publication and online: https://annovar.openbioinformatics.org/en/latest/user-guide/gene/. We scored all effect and non-effect alleles with each set of CNN models, **averaged predictions across folds, and calibrated CNN scores that predict activity using the set of validation positive OCRs. We computed the ΔSNP probability effect by taking the difference between the effect allele and the non-effect allele. Most SNPs reported by GWAS are not expected to be the causal variant for a trait, so the distribution of ΔSNP probability can be used to define a null distribution. We compute the p-value that an allele has a non-zero ΔSNP probability by fitting a normal distribution of null ΔSNP probabilities. We correct for multiple testing using the method swfdr v1.12.0 to compute q-values to control for a false-discoveries conditioned on potentially informative covariates (Boca and Leek, 2018). Weighted FDR-correction methods, including swfdr, have been shown to be robust to uninformative covariates and increase power to detect real differences for informative covariates while controlling false discoveries (Korthauer et al., 2019). We conditioned the proportion of expected null p-values on the following covariates (Supplemental Figure 5E, step 4): the difference in GC content of the 501 surrounding the SNP compared to the average GC content of positive sequences used to train each model (GC content), the minor allele frequency (MAF) based on the European ancestry subjects in the 1000G reference panel, whether the SNP overlapped a Fullard et al. NeuN+ OCR (inNeuN peak), and whether a SNP was fine-mapped and predicted to be causal by CAUSALdb using the European LD structure and an ensemble of statistical fine-mapping tools (isCausal) (FINEMAP, CAVIARBF, PAINTOR) (Chen et al., 2015; Benner et al., 2016; Kichaev et al., 2017; Wang et al., 2020). We applied an alpha of 0.05 on the false-discovery q-values for all 14,790 SNPs scored across 5 sets of CNNs to determine significantly large enough ΔSNP effects.**

To accompany cell type-specific activity predictions, we downloaded SNPs that are reported *cis* expression quantitative trait loci (eQTL) in human cortex, frontal cortex (DLPFC), anterior cingulate cortex (ACC), caudate, putamen, and the nucleus accumbens (NAc) from the GTEX Consortium from https://www.gtexportal.org/home/datasets(GTEx Consortium, 2013, 2015). We identified genes for which at least one of the 170 SNPs is an eQTL and plotted them as arcs in **Figure 4B** and **Supplemental Figure 4**. Locus plots are generated with pyGenomeTracks v3.5 tools (Ramírez et al., 2018).

For **Figure 4A**, we compared calibrated SNP probabilities of the either effect or non-effect allele across each model and grouped them by whether they overlapped a cortical or striatal NeuN+ OCR, NeuN-OCR, both, or neither, depending on whether the model was for EXC or D1/D2 neuronal subtypes, respectively. We computed 2-tailed t-tests between groups and corrected for multiple comparisons with the family-wise Bonferroni method for N=18 tests from three models and (4 choose 2) six possible comparisons per model. * P < 0.05/N, ** P < 0.01/N, *** P < 0.001/N.

## DATA AVAILABILITY

Code used to run intermediate and final analyses reported in this paper are available on the GitHub page: https://github.com/pfenninglab/addiction_gwas_enrichment. Sequencing output files for data generated in this study are deposited on the GEO at accession **GSE161374** (Reviewer access token: **cropkwsgnnyxhgh**). Questions and comments about data and analyses may be directed to the corresponding author.

## ACKNOWLEDGEMENT

We would like to thank members of the Eric Yttri lab at Carnegie Mellon University for providing *Drd1-cre* and *Adora2a-cre* mice for cell type-specific ATAC-seq experiments.

## Author contributions

Conceptualization: ARP, BNP, CS; ATAC-seq data processing: AJL, ER, IMK, BNP; GWAS enrichment investigation: BNP, CS, ER, MK; Machine learning models: AJL, BNP, ER, IMK; bulk tissue ATAC-seq: AJL, ARB, MEW; cSNAIL ATAC-seq: AJL, ARB; writing (original draft): BNP, CS; review and editing: AJL, ARP, BNP, CS, ER, IMK; funding acquisition, resources, & supervision: ARP;

---

**Supplemental Figure 1.**
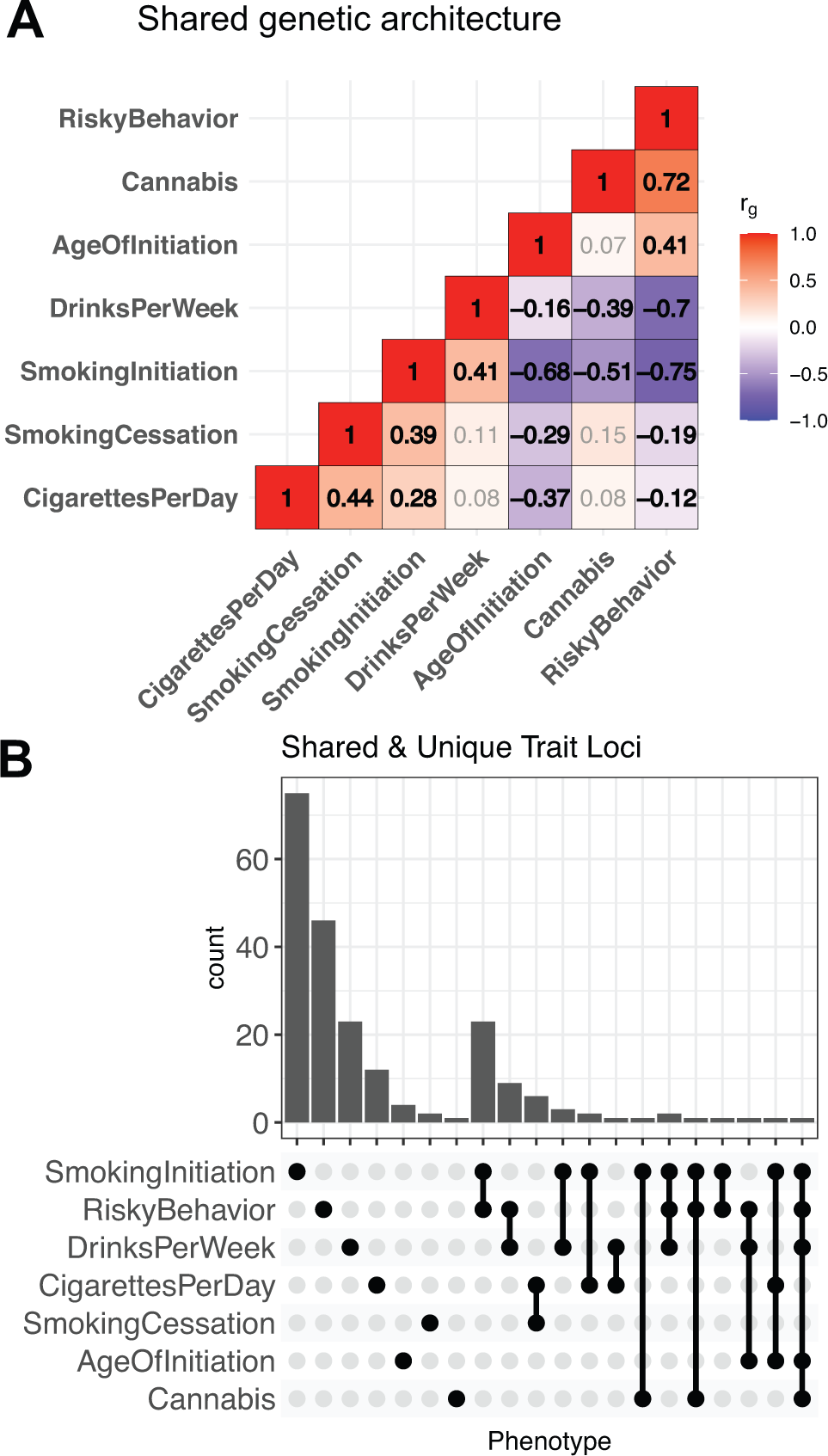
Shared and unique genetic architecture of genetic risk variants of addiction-associated traits. **(A)** LDSC genetic correlation (r_g_) matrix of seven addiction-associated traits. FDR-significant correlations at shown in bold, non-significant in gray (FDR < 0.05). **(B)** Upset plot of **non-overlapping** genomic loci shared or unique to each addiction-associated trait. Genomic loci are **clustered and identified** by shared GWAS-significant SNPs and genomic region overlap. **Supplemental Figure** . **Brain regions reported to be significantly enriched (FDR <= 0.05) are plotted with bolded bars, and the significance threshold is represented by a dashed red line. (C) Barplot of GWAS enrichment false-discovery rates in single cell open chromatin profiles of cell clusters in isocortex, hippocampus, and striatum** (Corces et al., 2020)**. Cell types in brain regions that are significantly enriched (FDR <= 0.05) are plotted with bolded bars, and the significance threshold is represented by a dashed red line. (D) Barplot of** GWAS enrichment false-discovery rates in single cell THS-seq OCRs of major cell clusters in occipital cortex (Lake et al., 2018). Cell types in brain regions that are significantly enriched (FDR <= 0.05) are plotted with **bolded bars, and the significance threshold is represented by a dashed red line**. Traits assessed are age of smoking initiation (AgeofInitiation), average number of cigarettes per day for ever smokers (CigarettesPerDay), having ever regularly smoked (SmokingInitiation), current versus former smokers (SmokingCessation), number of alcoholic drinks per week (DrinksPerWeek) (Liu et al., 2019b), lifetime cannabis use (Cannabis) (Pasman et al., 2018), and risky behavior (RiskyBehavior) (Karlsson Linnér et al., 2019). OFC: orbitofrontal cortex, VLPFC: ventrolateral prefrontal cortex, DLPFC: dorsolateral prefrontal cortex, ACC: anterior cingulate cortex, INS: insula, STC: superior temporal gyrus, ITC: inferior temporal gyrus, PMC: primary motor cortex, PVC: primary visual cortex, AMY: amygdala, HIPP: hippocampus, MDT: mediodorsal thalamus, NAc: nucleus accumbens, PUT: putamen, Ast: astrocyte, End: endothelial, Ex: excitatory neuron, In: inhibitory neuron, Mic: microglia, Oli: oligodendrocyte, Opc: oligodendrocyte precursor.

**Supplemental Figure 2.**
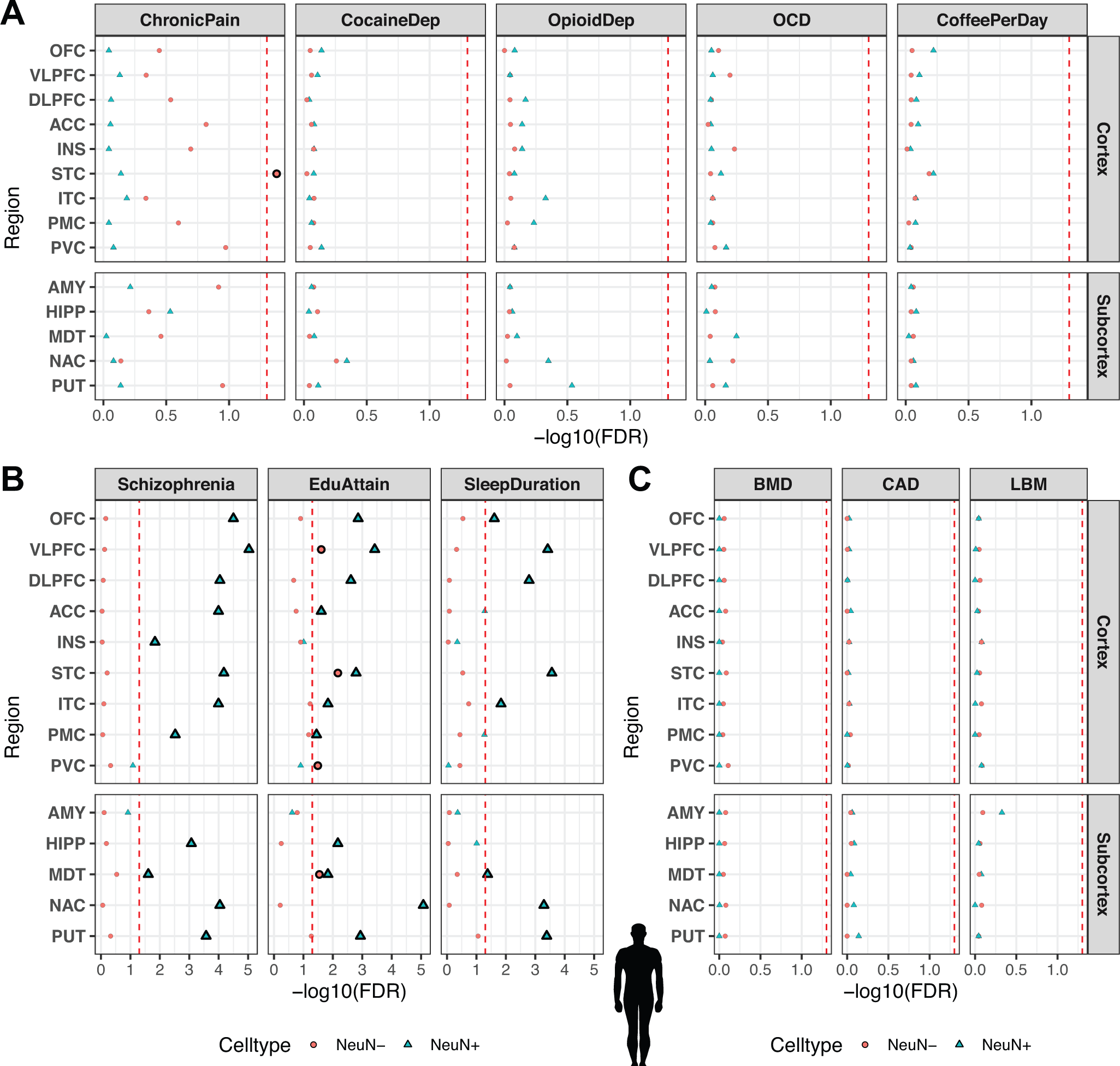
Sensitivity of partitioned LDSC regression for cell type- and region-specific in the GWAS trait enrichment requires well-powered GWAS in relevant cell types. GWAS enrichment plots with false-discovery rates in ATAC-seq of 14 postmortem human brain regions coupled with NeuN-labeled fluorescence activated nuclei sorting(Fullard et al., 2018). Regions are stratified by cortical and subcortical regions, with cortical regions ordered frontal to caudal. Sorted cell types within each brain region are denoted by shape (blue triangle for NeuN+/neuronal, red circle for NeuN-/glial). Cell types in brain regions that are enriched at FDR < 0.05 are plotted with bigger shapes and with black outlines. **(A)** GWAS enrichment of addiction-or substance use-associated traits: multi-site chronic pain (ChronicPain) (Johnston et al., 2019), cocaine dependence (CocaineDep) (Cabana-Domínguez et al., 2019), opioid dependence (OpioidDep)(Cheng et al., 2018), diagnosis of obsessive-compulsive disorder (OCD) (International Obsessive Compulsive Disorder Foundation Genetics Collaborative (IOCDF-GC) and OCD Collaborative Genetics Association Studies (OCGAS), 2018), and cups of coffee drank per day (CoffeePerDay) (Coffee and Caffeine Genetics Consortium et al., 2015). The GWAS for OCD, opioid dependence, and cocaine dependence are reportedly underpowered to detect genetic liability for these traits (N_case_< 5,000). **(B)** GWAS enrichment in well-powered brain-related traits show cell type- and region-specific enrichment: educational attainment (EduAttain) (Lee et al., 2018), schizophrenia risk (Schizophrenia) (Schizophrenia Working Group of the Psychiatric Genomics Consortium, 2014), habitual sleep duration (SleepDuration) (Dashti et al., 2019). **(C)** GWAS enrichment in non-brain associated traits do not show cell type- or region-specific enrichment: heel bone-mineral density (BMD) (Kemp et al., 2017), coronary artery disease (CAD) (Howson et al., 2017), and lean body mass (LBM) (Zillikens et al., 2017).

**Supplemental Figure 3.**
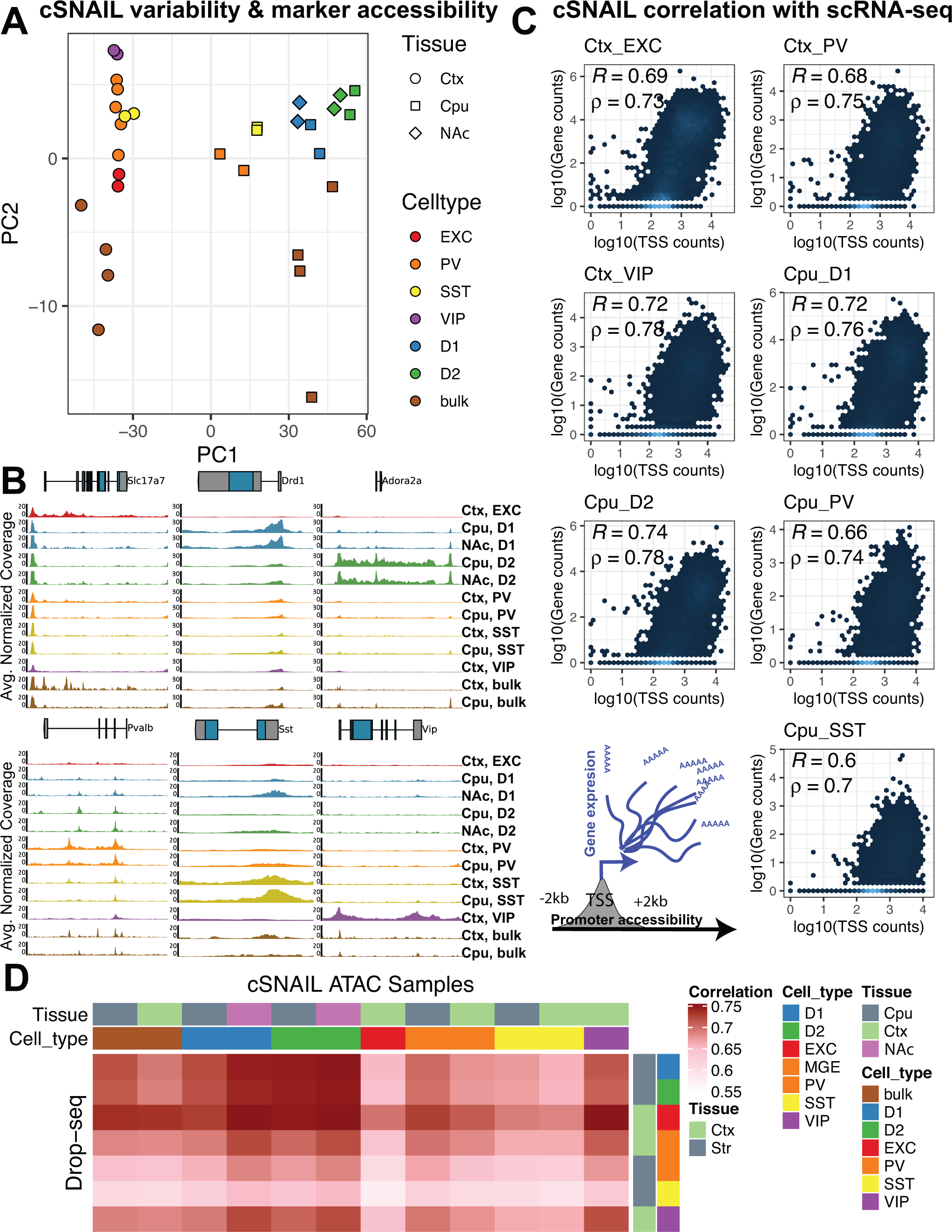
Cell type specificity of cSNAIL ATAC-seq in mouse cortex and striatum. (A) Principle component plots of chromatin accessibility counts from *cre*-dependent Sun1-GFP Nuclear Anchored Independent Labeled (cSNAIL) ATAC-seq from *cre*-driver lines (**Methods**). Major axes of variation separate cell types by tissue source (PC1) and cell type versus bulk ATAC-seq (PC2). (B) Normalized coverage track plots around marker genes demarcating cell type-specificity of cSNAIL ATAC-seq samples. (C) Density correlation plot of normalized chromatin accessibility log counts around the transcription start site (TSS) **correlated** with matched pseudo-bulk cell type log gene counts from Drop-seq of mouse cortex and striatum(Saunders et al., 2018). Drop-seq cell types meta-gene profiles report sum gene counts for cell clusters from frontal cortex and striatum. Pearson’s and Spearman’s correlation are denoted with R and ρ, respectively. (D) Pairwise correlation matrix of TSS chromatin accessibility log counts with Drop-seq pseudo-bulk log gene counts from cortical and striatal cell clusters.

**Supplemental Figure 4.**
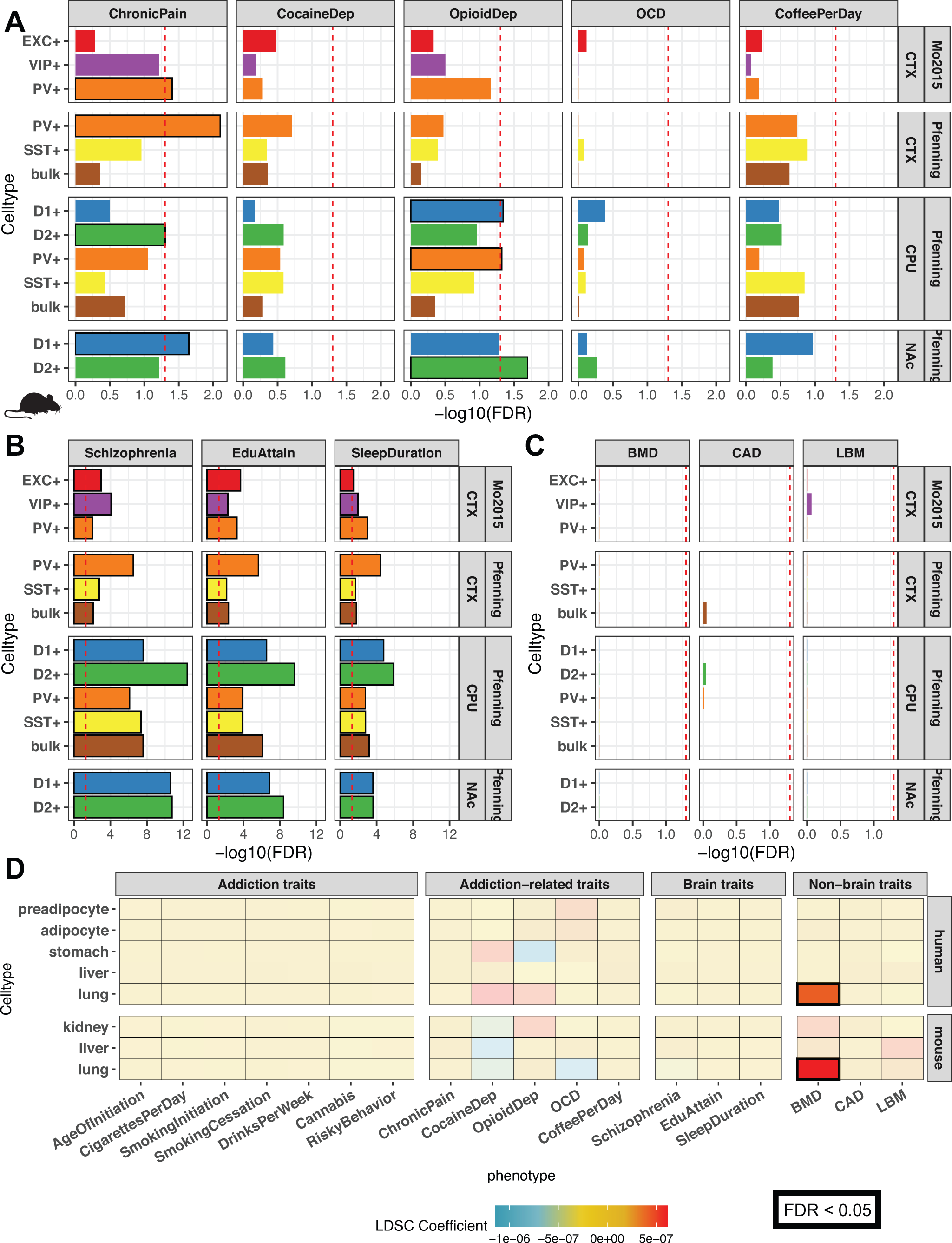
GWAS enrichment in addiction- and non-addiction-related traits using mapped mouse orthologs of tissue- and cell type-specific open chromatin regions. GWAS enrichment plots with false-discovery rates in human orthologous regions mapped from mouse ATAC-seq of bulk cortex (CTX), dorsal striatum (CPU), and nucleus accumbens (**NAc**) or cre-dependent Sun1-GFP Nuclear Anchored Independent Labeled (cSNAIL) nuclei of D1-cre, D2-cre, and PV-cre mice. cSNAIL ATAC-seq experiments report both enriched (+) and de-enriched (-) nuclei populations. Enrichments that are enriched at FDR < 0.05 are plot with black outlines. Replication of enrichment is shown using INTACT-enriched OCRs from Mo *et al*(Mo et al., 2015) of cortical excitatory (EXC+), vasoactive intestinal peptide interneuron (VIP+), and parvalbumin interneuron (PV+). **(A)** GWAS enrichment of addiction- or substance use-associated traits: multi-site chronic pain (ChronicPain), cocaine dependence (CocaineDep), opioid dependence (OpioidDep), diagnosis of obsessive-compulsive disorder (OCD), and cups of coffee drank per day (CoffeePerDay). The GWAS for OCD, opioid dependence, and cocaine dependence are reportedly underpowered to detect genetic liability for these traits (N_case_< 5,000). **(B)** GWAS enrichment in well-powered brain-related traits show cell type- and region-specific enrichment: educational attainment (EduAttain), schizophrenia risk (Schizophrenia), habitual sleep duration (SleepDuration). **(C)** GWAS enrichment in non-brain associated traits do not show cell type- or region-specific enrichment: heel bone-mineral density (BMD), coronary artery disease (CAD), and lean body mass (LBM). **(D)** Heatmap of LDSC regression coefficients of GWAS enrichment for all measured GWAS in non-brain OCRs from human or mouse-human mapped orthologs. Tissues for which OCRs are significantly enriched (FDR < 0.05) with GWAS variants are outlined with a bolded box.

**Supplemental Figure 5.**
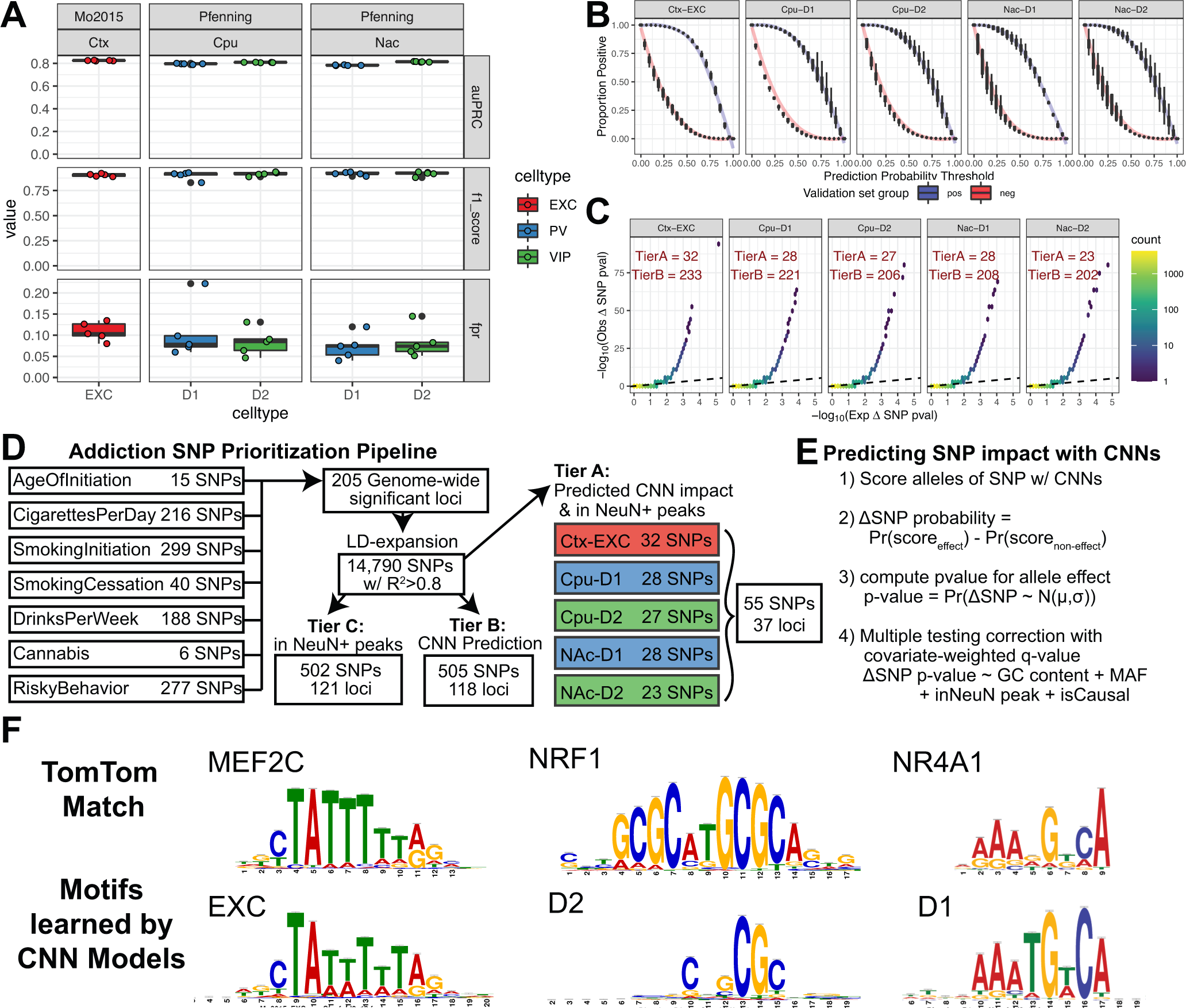
Convolutional Neural Network (CNN) model performance and selection of candidate functional SNPs. **(A)** Performance metrics for **convolutional neural network (CNN)** models **show high specificity on the test sets of positive peaks or 10x nucleotide-content matched negatives. Test set** performance metrics are reported for area under the precision-recall curve (auPRC)**, F1-score (using threshold = 0.5), and false positive rates across all possible thresholds** (**Methods**). Models were trained on IDR peaks of mouse cortical excitatory **(Ctx-EXC)** and D1 and D2 medium spiny **neurons from caudoputamen (CPU) and the nucleus accumbens (NAc). (B) The models best discriminate the proportion of positives and negative sequences at a threshold of 0.5. Plots show the proportion of positives (blue) or negatives (red) that are called “positive” across CNN output thresholds from 0 to 1 averaged across folds for each set of CNN models. (C) Quantile-quantile plots of p-values of calibrated ΔSNP probability (Methods) from a normal distribution after centering by the mean and scaling by the standard deviation of delta SNP probabilities across all SNPs (n=14,790 SNPs) for each set of CNN models. A hexbin plot was used instead to better visualize over-plotting where every hexagon is color by the number of SNPs in that observed and expected p-value. The black dotted line denotes the equality line y = x. The number of significant SNPs at false discovery q-value < 0.05 at Tier A or B are reported for each cell type and tissue (Methods). (D) Schematic to select for predicted causal impact addiction-associated GWAS SNPs. The pipeline begins with SNPs across addiction-associated GWAS aggregated to 205 non-overlapping GWAS loci across 14,790 SNPs after LD-expansion to include those in LD R^2^ > 0.8. SNPs are further prioritized into three tiers. Tier C includes SNPs with only overlap with Fullard *et al*. NeuN+ ATAC-seq peaks, Tier B includes SNPs with only predicted significant differential allelic impact by on CNN-predicted CRE activity at q-value < 0.05, and Tier A include SNPs matching both criteria (Methods). (E) Outline of predicting differential CRE activity between alleles using calibrated CNN probabilities of CRE activity while controlling for false discovery using informative covariates (Methods). (F) Example motif matches from Supplemental Table 2 of TomTom known transcription factor consensus motifs and the learned important features in CNN models for cortical excitatory and striatal D1 and D2 MSNs.**

**Supplemental Figure 6.**
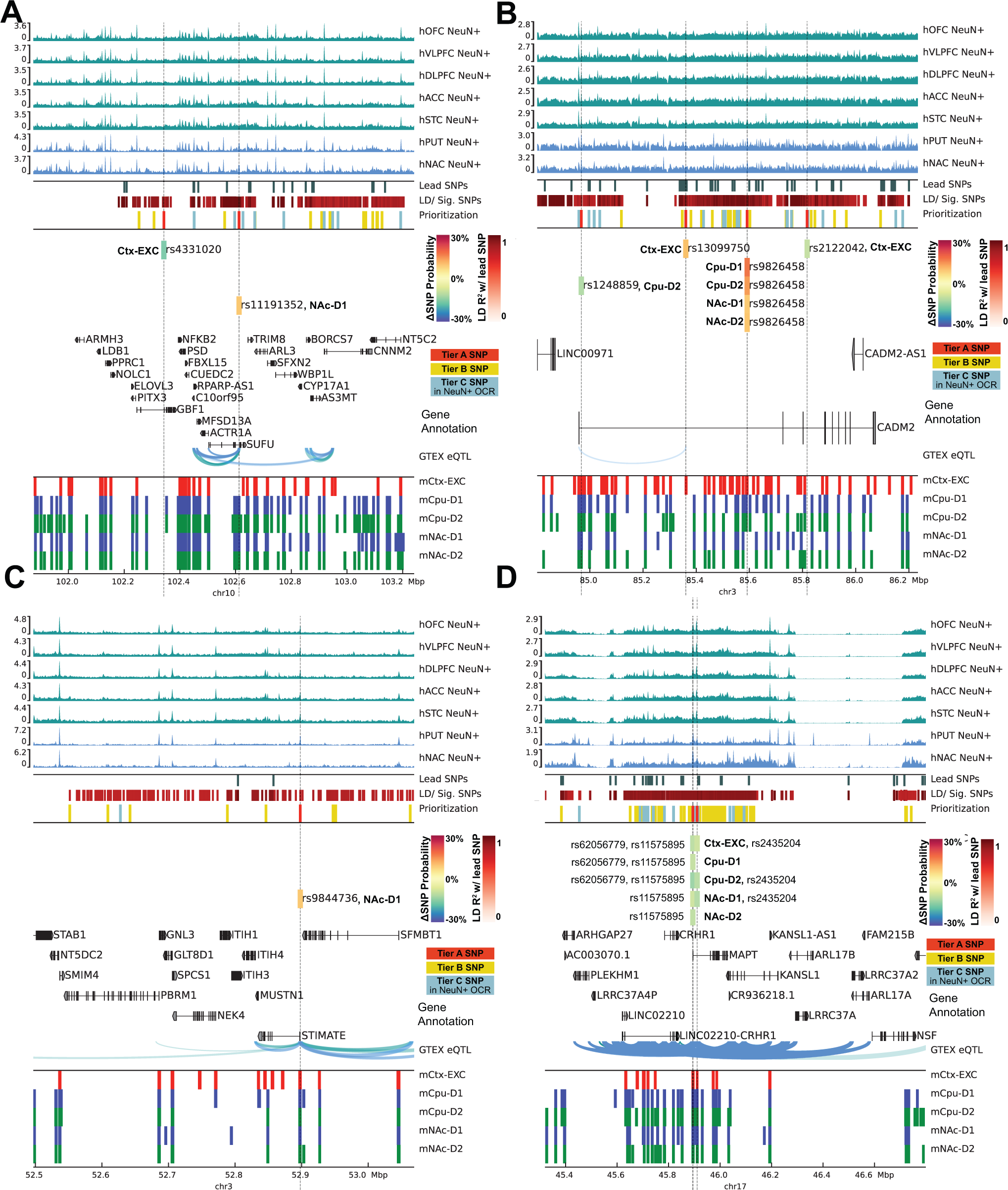
Locus plots of addiction-associated SNPs predicted to act in striatal and cortical cell types. Locus plot across four additional loci with Tier A SNPs with predicted function SNP impact in cortical excitatory and striatal D1 and D2 MSN cell types. Genome tracks from top to bottom: human (h)NeuN+ MACS2 ATAC-seq fold change signal of cortical and striatal brain regions enriched in Figure 1A. SNP tracks plot lead SNPs aggregated across seven addiction-associated GWAS, the SNPs in LD with the lead SNPs (Lead SNPs) or independently significant SNPs (LD/ Sig. SNPs). Each SNP is colored by red increasing in intensity by the degree of LD with a lead SNP. Prioritized candidate causal SNPs by predicted differential cell type activity and overlap with Fullard *et al*. NeuN+ OCRs are plot as (red for Tier A, yellow for Tier B, and teal for Tier C, Methods). Tier A SNP rs7604640 is predicted to have strong ΔSNP effect by CPU-D1 and NAc-D1 CNN models and the bars are colored by the % change in probability active. Gene annotation tracks plot GENCODE genes from the GRCh38 build. eQTL link tracks of FDR-significant GTEX cis-eQTL from cortical and striatal brain regions, and orthologs of mouse (m) putative CREs mapped from excitatory or striatal neuronal subtypes measured by cSNAIL ATAC-seq. NeuN+ ATAC-seq tracks and eQTL links are colored by source brain region as cortical (teal) or striatal (blue). Cell type colors label cortical excitatory neurons (EXC; red), D1 medium spiny neurons (D1; blue), or D2 medium spiny neurons (D2; green).

**Supplemental Table 1. Addiction-associated genetic variants annotated with cell type and brain region functional markers**

**Addiction-associated genetic variants from the main seven GWAS (Figure 1) that were scored by CNN models along with computed raw CNN scores, predicted probability active, and ΔSNP probabilities, and tier of predicted candidate causal SNP. Each entry is recorded for a distinct SNP, predicted CNN model, and GWAS trait. Additional columns reporting are** annotated by FUMA (Watanabe et al., 2017) **and** CAUSALdb **(Wang et al., 2020). SNPs are annotated in this study to** overlap with **human** NeuN+ OCRs (Fullard et al., 2018). **A complete legend describing column headers is in the first sheet of the table.**

**Supplemental Table 2. TomTom matches with motifs learned by CNN models in each cell type and fold to contribute to a strong positive prediction. Learned important features were interpreted by DeepSHAP and clustered into unique seqlets by TF-Modisco (Methods).**

## Bibliography

1000 Genomes Project Consortium, Auton A, Brooks LD, Durbin RM, Garrison EP, Kang HM, Korbel JO, Marchini JL, McCarthy S, McVean GA, Abecasis GR (2015) A global reference for human genetic variation. Nature 526:68–74.

Alipanahi B, Delong A, Weirauch MT, Frey BJ (2015) Predicting the sequence specificities of DNA- and RNA-binding proteins by deep learning. Nat Biotechnol 33:831–838.

Barman P, Reddy D, Bhaumik SR (2019) Mechanisms of Antisense Transcription Initiation with Implications in Gene Expression, Genomic Integrity and Disease Pathogenesis. Non-coding RNA 5.

Beaulieu C (1993) Numerical data on neocortical neurons in adult rat, with special reference to the GABA population. Brain Res 609:284–292.

Benner C, Spencer CCA, Havulinna AS, Salomaa V, Ripatti S, Pirinen M (2016) FINEMAP: efficient variable selection using summary data from genome-wide association studies. Bioinformatics 32:1493–1501.

Berke JD, Hyman SE (2000) Addiction, dopamine, and the molecular mechanisms of memory. Neuron 25:515–532.

Boca SM, Leek JT (2018) A direct approach to estimating false discovery rates conditional on covariates. PeerJ 6:e6035.

Bracci E, Centonze D, Bernardi G, Calabresi P (2002) Dopamine excites fast-spiking interneurons in the striatum. J Neurophysiol 87:2190–2194.

Buenrostro JD, Giresi PG, Zaba LC, Chang HY, Greenleaf WJ (2013) Transposition of native chromatin for fast and sensitive epigenomic profiling of open chromatin, DNA-binding proteins and nucleosome position. Nat Methods 10:1213–1218.

Buenrostro JD, Wu B, Chang HY, Greenleaf WJ (2015) ATAC-seq: A Method for Assaying Chromatin Accessibility Genome-Wide. Curr Protoc Mol Biol 109:21.29.1-21.29.9.

Bulik-Sullivan B, Finucane HK, Anttila V, Gusev A, Day FR, Loh P-R, ReproGen Consortium, Psychiatric Genomics Consortium, Genetic Consortium for Anorexia Nervosa of the Wellcome Trust Case Control Consortium 3, Duncan L, Perry JRB, Patterson N, Robinson EB, Daly MJ, Price AL, Neale BM (2015a) An atlas of genetic correlations across human diseases and traits. Nat Genet 47:1236–1241.

Bulik-Sullivan B, Loh P-R, Finucane HK, Ripke S, Yang J, Schizophrenia Working Group of the Psychiatric Genomics Consortium, Patterson N, Daly MJ, Price AL, Neale BM (2015b) LD Score regression distinguishes confounding from polygenicity in genome-wide association studies. Nat Genet 47:291–295.

Bush WS, Moore JH (2012) Chapter 11: Genome-wide association studies. PLoS Comput Biol 8:e1002822.

Cabana-Domínguez J, Shivalikanjli A, Fernàndez-Castillo N, Cormand B (2019) Genome-wide association meta-analysis of cocaine dependence: Shared genetics with comorbid conditions. Prog Neuropsychopharmacol Biol Psychiatry 94:109667.

Cannon ME, Currin KW, Young KL, Perrin HJ, Vadlamudi S, Safi A, Song L, Wu Y, Wabitsch M, Laakso M, Crawford GE, Mohlke KL (2019) Open Chromatin Profiling in Adipose Tissue Marks Genomic Regions with Functional Roles in Cardiometabolic Traits. G3 (Bethesda) 9:2521–2533.

Chan KY, Jang MJ, Yoo BB, Greenbaum A, Ravi N, Wu W-L, Sánchez-Guardado L, Lois C, Mazmanian SK, Deverman BE, Gradinaru V (2017) Engineered AAVs for efficient noninvasive gene delivery to the central and peripheral nervous systems. Nat Neurosci 20:1172–1179.

Chen L, Fish AE, Capra JA (2018) Prediction of gene regulatory enhancers across species reveals evolutionarily conserved sequence properties. PLoS Comput Biol 14:e1006484.

Chen W, Larrabee BR, Ovsyannikova IG, Kennedy RB, Haralambieva IH, Poland GA, Schaid DJ (2015) Fine Mapping Causal Variants with an Approximate Bayesian Method Using Marginal Test Statistics. Genetics 200:719–736.

Cheng Y, Huang CCY, Ma T, Wei X, Wang X, Lu J, Wang J (2017) Distinct synaptic strengthening of the striatal direct and indirect pathways drives alcohol consumption. Biol Psychiatry 81:918–929.

Cheng Z, Zhou H, Sherva R, Farrer LA, Kranzler HR, Gelernter J (2018) Genome-wide Association Study Identifies a Regulatory Variant of RGMA Associated With Opioid Dependence in European Americans. Biol Psychiatry 84:762–770.

Coffee and Caffeine Genetics Consortium et al. (2015) Genome-wide meta-analysis identifies six novel loci associated with habitual coffee consumption. Mol Psychiatry 20:647–656.

Corces MR, Shcherbina A, Kundu S, Gloudemans MJ, Frésard L, Granja JM, Louie BH, Eulalio T, Shams S, Bagdatli ST, Mumbach MR, Liu B, Montine KS, Greenleaf WJ, Kundaje A, Montgomery SB, Chang HY, Montine TJ (2020) Single-cell epigenomic analyses implicate candidate causal variants at inherited risk loci for Alzheimer’s and Parkinson’s diseases. Nat Genet 52:1158–1168.

Dashti HS et al. (2019) Genome-wide association study identifies genetic loci for self-reported habitual sleep duration supported by accelerometer-derived estimates. Nat Commun 10:1100.

Davis CA, Hitz BC, Sloan CA, Chan ET, Davidson JM, Gabdank I, Hilton JA, Jain K, Baymuradov UK, Narayanan AK, Onate KC, Graham K, Miyasato SR, Dreszer TR, Strattan JS, Jolanki O, Tanaka FY, Cherry JM (2018) The Encyclopedia of DNA elements (ENCODE): data portal update. Nucleic Acids Res 46:D794–D801.

Dick DM (2016) The genetics of addiction: where do we go from here? J Stud Alcohol Drugs 77:673–675.

Eddie D, Greene MC, White WL, Kelly JF (2019) Medical burden of disease among individuals in recovery from alcohol and other drug problems in the united states: findings from the national recovery survey. J Addict Med 13:385–395.

ENCODE Project Consortium (2012) An integrated encyclopedia of DNA elements in the human genome. Nature 489:57–74.

Erzurumluoglu AM et al. (2019) Meta-analysis of up to 622,409 individuals identifies 40 novel smoking behaviour associated genetic loci. Mol Psychiatry.

Farrell MR, Schoch H, Mahler SV (2018) Modeling cocaine relapse in rodents: Behavioral considerations and circuit mechanisms. Prog Neuropsychopharmacol Biol Psychiatry 87:33–47.

Fehr C, Yakushev I, Hohmann N, Buchholz H-G, Landvogt C, Deckers H, Eberhardt A, Kläger M, Smolka MN, Scheurich A, Dielentheis T, Schmidt LG, Rösch F, Bartenstein P, Gründer G, Schreckenberger M (2008) Association of low striatal dopamine d2 receptor availability with nicotine dependence similar to that seen with other drugs of abuse. Am J Psychiatry 165:507–514.

Ferguson SM, Eskenazi D, Ishikawa M, Wanat MJ, Phillips PEM, Dong Y, Roth BL, Neumaier JF (2011) Transient neuronal inhibition reveals opposing roles of indirect and direct pathways in sensitization. Nat Neurosci 14:22–24.

Finucane HK et al. (2015) Partitioning heritability by functional annotation using genome-wide association summary statistics. Nat Genet 47:1228–1235.

Finucane HK et al. (2018) Heritability enrichment of specifically expressed genes identifies disease-relevant tissues and cell types. Nat Genet 50:621–629.

Fullard JF, Hauberg ME, Bendl J, Egervari G, Cirnaru M-D, Reach SM, Motl J, Ehrlich ME, Hurd YL, Roussos P (2018) An atlas of chromatin accessibility in the adult human brain. Genome Res 28:1243–1252.

GBD 2016 Alcohol and Drug Use Collaborators (2018) The global burden of disease attributable to alcohol and drug use in 195 countries and territories, 1990-2016: a systematic analysis for the Global Burden of Disease Study 2016. Lancet Psychiatry 5:987–1012.

Ghandi M, Lee D, Mohammad-Noori M, Beer MA (2014) Enhanced regulatory sequence prediction using gapped k-mer features. PLoS Comput Biol 10:e1003711.

Gjoneska E, Pfenning AR, Mathys H, Quon G, Kundaje A, Tsai L-H, Kellis M (2015) Conserved epigenomic signals in mice and humans reveal immune basis of Alzheimer’s disease. Nature 518:365–369.

Goldstein RZ, Volkow ND (2011) Dysfunction of the prefrontal cortex in addiction: neuroimaging findings and clinical implications. Nat Rev Neurosci 12:652–669.

Grillner S, Robertson B (2016) The basal ganglia over 500 million years. Curr Biol 26:R1088– R1100.

GTEx Consortium (2013) The Genotype-Tissue Expression (GTEx) project. Nat Genet 45:580– 585.

GTEx Consortium (2015) Human genomics. The Genotype-Tissue Expression (GTEx) pilot analysis: multitissue gene regulation in humans. Science 348:648–660.

GTEx Consortium et al. (2017) Genetic effects on gene expression across human tissues. Nature 550:204–213.

Gupta S, Stamatoyannopoulos JA, Bailey TL, Noble WS (2007) Quantifying similarity between motifs. Genome Biol 8:R24.

Hickey G, Paten B, Earl D, Zerbino D, Haussler D (2013) HAL: a hierarchical format for storing and analyzing multiple genome alignments. Bioinformatics 29:1341–1342.

Hodge RD et al. (2019) Conserved cell types with divergent features in human versus mouse cortex. Nature 573:61–68.

Hoffman JL, Faccidomo S, Kim M, Taylor SM, Agoglia AE, May AM, Smith EN, Wong LC, Hodge CW (2019) Alcohol Drinking Exacerbates Neural and Behavioral Pathology in the 3xTg-AD Mouse Model of Alzheimer’s Disease. BioRxiv.

Howson JMM et al. (2017) Fifteen new risk loci for coronary artery disease highlight arterial-wall-specific mechanisms. Nat Genet 49:1113–1119.

International Obsessive Compulsive Disorder Foundation Genetics Collaborative (IOCDF-GC) and OCD Collaborative Genetics Association Studies (OCGAS) (2018) Revealing the complex genetic architecture of obsessive-compulsive disorder using meta-analysis. Mol Psychiatry 23:1181–1188.

Jensen KP (2016) A Review of Genome-Wide Association Studies of Stimulant and Opioid Use Disorders. Mol Neuropsychiatry 2:37–45.

Ji X, Saha S, Kolpakova J, Guildford M, Tapper AR, Martin GE (2017) Dopamine Receptors Differentially Control Binge Alcohol Drinking-Mediated Synaptic Plasticity of the Core Nucleus Accumbens Direct and Indirect Pathways. J Neurosci 37:5463–5474.

Jiang C, Wang X, Le Q, Liu P, Liu C, Wang Z, He G, Zheng P, Wang F, Ma L (2019) Morphine coordinates SST and PV interneurons in the prelimbic cortex to disinhibit pyramidal neurons and enhance reward. Mol Psychiatry.

Johnston KJA, Adams MJ, Nicholl BI, Ward J, Strawbridge RJ, Ferguson A, McIntosh AM, Bailey MES, Smith DJ (2019) Genome-wide association study of multisite chronic pain in UK Biobank. PLoS Genet 15:e1008164.

Karlsson Linnér R et al. (2019) Genome-wide association analyses of risk tolerance and risky behaviors in over 1 million individuals identify hundreds of loci and shared genetic influences. Nat Genet 51:245–257.

Kelley DR, Reshef YA, Bileschi M, Belanger D, McLean CY, Snoek J (2018) Sequential regulatory activity prediction across chromosomes with convolutional neural networks. Genome Res 28:739–750.

Kelley DR, Snoek J, Rinn JL (2016) Basset: learning the regulatory code of the accessible genome with deep convolutional neural networks. Genome Res 26:990–999.

Kemp JP et al. (2017) Identification of 153 new loci associated with heel bone mineral density and functional involvement of GPC6 in osteoporosis. Nat Genet 49:1468–1475.

Kendler KS, Prescott CA (1998a) Cannabis use, abuse, and dependence in a population-based sample of female twins. Am J Psychiatry 155:1016–1022.

Kendler KS, Prescott CA (1998b) Cocaine use, abuse and dependence in a population-based sample of female twins. Br J Psychiatry 173:345–350.

Khan A, Riudavets Puig R, Boddie P, Mathelier A (2020) BiasAway: command-line and web server to generate nucleotide composition-matched DNA background sequences. Available at: https://biasaway.uio.no [Accessed July 1, 2020].

Kichaev G, Roytman M, Johnson R, Eskin E, Lindström S, Kraft P, Pasaniuc B (2017) Improved methods for multi-trait fine mapping of pleiotropic risk loci. Bioinformatics 33:248–255.

Kim C, Kim S, Lee KY, Kim NH, Kang E-Y, Oh Y-W, Shin C (2019) Differences in bone density on chest CT according to smoking status in males without chronic obstructive lung disease. Sci Rep 9:10467.

Koob GF, Volkow ND (2010) Neurocircuitry of addiction. Neuropsychopharmacology 35:217– 238.

Koob GF, Volkow ND (2016) Neurobiology of addiction: a neurocircuitry analysis. Lancet Psychiatry 3:760–773.

Korthauer K, Kimes PK, Duvallet C, Reyes A, Subramanian A, Teng M, Shukla C, Alm EJ, Hicks SC (2019) A practical guide to methods controlling false discoveries in computational biology. Genome Biol 20:118.

Krienen FM et al. (2019) Innovations in primate interneuron repertoire. BioRxiv.

Lake BB, Chen S, Sos BC, Fan J, Kaeser GE, Yung YC, Duong TE, Gao D, Chun J, Kharchenko PV, Zhang K (2018) Integrative single-cell analysis of transcriptional and epigenetic states in the human adult brain. Nat Biotechnol 36:70–80.

Landt SG et al. (2012) ChIP-seq guidelines and practices of the ENCODE and modENCODE consortia. Genome Res 22:1813–1831.

Lansink CS, Goltstein PM, Lankelma JV, Pennartz CMA (2010) Fast-spiking interneurons of the rat ventral striatum: temporal coordination of activity with principal cells and responsiveness to reward. Eur J Neurosci 32:494–508.

Lawler AJ, Brown AR, Bouchard RS, Toong N, Kim Y, Velraj N, Fox G, Kleyman M, Kang B, Gittis AH, Pfenning AR (2020) Cell Type-Specific Oxidative Stress Genomic Signatures in the Globus Pallidus of Dopamine-Depleted Mice. J Neurosci 40:9772–9783.

Lee D (2016) LS-GKM: a new gkm-SVM for large-scale datasets. Bioinformatics 32:2196– 2198.

Lee IS, Leem AY, Lee SH, Rhee Y, Ha Y, Kim YS (2016) Relationship between pulmonary function and bone mineral density in the Korean National Health and Nutrition Examination Survey. Korean J Intern Med 31:899–909.

Lee JH, Ribeiro EA, Kim J, Ko B, Kronman H, Jeong YH, Kim JK, Janak PH, Nestler EJ, Koo JW, Kim J-H (2020) Dopaminergic regulation of nucleus accumbens cholinergic interneurons demarcates susceptibility to cocaine addiction. Biol Psychiatry.

Lee JJ et al. (2018) Gene discovery and polygenic prediction from a genome-wide association study of educational attainment in 1.1 million individuals. Nat Genet 50:1112–1121.

Lefort S, Tomm C, Floyd Sarria JC, Petersen CCH (2009) The excitatory neuronal network of the C2 barrel column in mouse primary somatosensory cortex. Neuron 61:301–316.

Liao Y, Smyth GK, Shi W (2014) featureCounts: an efficient general purpose program for assigning sequence reads to genomic features. Bioinformatics 30:923–930.

Liu C, Wang M, Wei X, Wu L, Xu J, Dai X, Xia J, Cheng M, Yuan Y, Zhang P, Li J, Feng T, Chen A, Zhang W, Chen F, Shang Z, Zhang X, Peters BA, Liu L (2019a) An ATAC-seq atlas of chromatin accessibility in mouse tissues. Sci Data 6:65.

Liu M et al. (2019b) Association studies of up to 1.2 million individuals yield new insights into the genetic etiology of tobacco and alcohol use. Nat Genet 51:237–244.

Love MI, Huber W, Anders S (2014) Moderated estimation of fold change and dispersion for RNA-seq data with DESeq2. Genome Biol 15:550–550.

Maurano MT et al. (2012) Systematic localization of common disease-associated variation in regulatory DNA. Science 337:1190–1195.

Melé M et al. (2015) The human transcriptome across tissues and individuals. Science 348:660– 665.

Minnoye L, Taskiran II, Mauduit D, Fazio M, Van Aerschot L, Hulselmans G, Christiaens V, Makhzami S, Seltenhammer M, Karras P, Primot A, Cadieu E, van Rooijen E, Marine J- C, Egidy G, Ghanem GE, Zon L, Wouters J, Aerts S (2020) Cross-species analysis of enhancer logic using deep learning. Genome Res.

Mo A, Mukamel EA, Davis FP, Luo C, Henry GL, Picard S, Urich MA, Nery JR, Sejnowski TJ, Lister R, Eddy SR, Ecker JR, Nathans J (2015) Epigenomic signatures of neuronal diversity in the mammalian brain. Neuron 86:1369–1384.

Monaco G, van Dam S, Casal Novo Ribeiro JL, Larbi A, de Magalhães JP (2015) A comparison of human and mouse gene co-expression networks reveals conservation and divergence at the tissue, pathway and disease levels. BMC Evol Biol 15:259.

Pasman JA et al. (2018) GWAS of lifetime cannabis use reveals new risk loci, genetic overlap with psychiatric traits, and a causal influence of schizophrenia. Nat Neurosci 21:1161– 1170.

Paten B, Earl D, Nguyen N, Diekhans M, Zerbino D, Haussler D (2011) Cactus: Algorithms for genome multiple sequence alignment. Genome Res 21:1512–1528.

Pear VA, Ponicki WR, Gaidus A, Keyes KM, Martins SS, Fink DS, Rivera-Aguirre A, Gruenewald PJ, Cerdá M (2019) Urban-rural variation in the socioeconomic determinants of opioid overdose. Drug Alcohol Depend 195:66–73.

Pelechano V, Steinmetz LM (2013) Gene regulation by antisense transcription. Nat Rev Genet 14:880–893.

Pullen E, Oser C (2014) Barriers to substance abuse treatment in rural and urban communities: counselor perspectives. Subst Use Misuse 49:891–901.

Purcell S, Neale B, Todd-Brown K, Thomas L, Ferreira MAR, Bender D, Maller J, Sklar P, de Bakker PIW, Daly MJ, Sham PC (2007) PLINK: a tool set for whole-genome association and population-based linkage analyses. Am J Hum Genet 81:559–575.

Ramamurthy E, Welch G, Cheng J, Yuan Y, Gunsalus L, Bennett DA, Tsai L-H, Pfenning A (2020) Cell type-specific histone acetylation profiling of Alzheimer’s Disease subjects and integration with genetics. BioRxiv.

Ramírez F, Bhardwaj V, Arrigoni L, Lam KC, Grüning BA, Villaveces J, Habermann B, Akhtar A, Manke T (2018) High-resolution TADs reveal DNA sequences underlying genome organization in flies. Nat Commun 9:189.

Ramírez F, Ryan DP, Grüning B, Bhardwaj V, Kilpert F, Richter AS, Heyne S, Dündar F, Manke T (2016) deepTools2: a next generation web server for deep-sequencing data analysis. Nucleic Acids Res 44:W160–5.

Ribeiro EA et al. (2018) Transcriptional and physiological adaptations in nucleus accumbens somatostatin interneurons that regulate behavioral responses to cocaine. Nat Commun 9:3149.

Roadmap Epigenomics Consortium et al. (2015) Integrative analysis of 111 reference human epigenomes. Nature 518:317–330.

Saunders A, Macosko EZ, Wysoker A, Goldman M, Krienen FM, de Rivera H, Bien E, Baum M, Bortolin L, Wang S, Goeva A, Nemesh J, Kamitaki N, Brumbaugh S, Kulp D, McCarroll SA (2018) Molecular Diversity and Specializations among the Cells of the Adult Mouse Brain. Cell 174:1015–1030.e16.

Scaplen KM, Kaun KR (2016) Reward from bugs to bipeds: a comparative approach to understanding how reward circuits function. J Neurogenet 30:133–148.

Schall TA, Wright WJ, Dong Y (2020) Nucleus accumbens fast-spiking interneurons in motivational and addictive behaviors. Mol Psychiatry.

Schizophrenia Working Group of the Psychiatric Genomics Consortium (2014) Biological insights from 108 schizophrenia-associated genetic loci. Nature 511:421–427.

Seney M, Kim S-M, Wang J, Hildebrand M, Xue X, Glausier J, Zong W, Shelton M, Phan B, Srinivasan C, Pfenning A, Tseng G, Lewis D, Freyberg Z, Logan R (2020) Transcriptional alterations in opioid use disorder reveal an interplay between neuroinflammation and synaptic remodeling. BioRxiv.

Shlyueva D, Stampfel G, Stark A (2014) Transcriptional enhancers: from properties to genome-wide predictions. Nat Rev Genet 15:272–286.

Shrikumar A, Greenside P, Kundaje A (2017) Learning Important Features Through Propagating Activation Differences. arXiv.

Shrikumar A, Tian K, Avsec Ž, Shcherbina A, Banerjee A, Sharmin M, Nair S, Kundaje A (2018) Technical Note on Transcription Factor Motif Discovery from Importance Scores (TF-MoDISco) version 0.5.6.5. arXiv.

Smith LN (2018) A disciplined approach to neural network hyper-parameters: Part 1 -- learning rate, batch size, momentum, and weight decay. arXiv.

Spitz F, Furlong EEM (2012) Transcription factors: from enhancer binding to developmental control. Nat Rev Genet 13:613–626.

Štrumbelj E, Kononenko I (2014) Explaining prediction models and individual predictions with feature contributions. Knowl Inf Syst 41:647–665.

Tak YG, Farnham PJ (2015) Making sense of GWAS: using epigenomics and genome engineering to understand the functional relevance of SNPs in non-coding regions of the human genome. Epigenetics Chromatin 8:57.

Tepper JM, Koós T (2017) Gabaergic interneurons of the striatum. In: Handbook of basal ganglia structure and function, second edition, pp 157–178 Handbook of behavioral neuroscience. Elsevier.

Thurman RE et al. (2012) The accessible chromatin landscape of the human genome. Nature 489:75–82.

Visscher PM, Wray NR, Zhang Q, Sklar P, McCarthy MI, Brown MA, Yang J (2017) 10 years of GWAS discovery: biology, function, and translation. Am J Hum Genet 101:5–22.

Volkow ND, Chang L, Wang G-J, Fowler JS, Ding Y-S, Sedler M, Logan J, Franceschi D, Gatley J, Hitzemann R, Gifford A, Wong C, Pappas N (2003) Low level of brain dopamine d_2_ receptors in methamphetamine abusers: association with metabolism in the orbitofrontal cortex. Focus (Madison) 1:150–157.

Volkow ND, Morales M (2015) The brain on drugs: from reward to addiction. Cell 162:712– 725.

Volkow ND, Wang GJ, Fowler JS, Logan J, Gatley SJ, Hitzemann R, Chen AD, Dewey SL, Pappas N (1997) Decreased striatal dopaminergic responsiveness in detoxified cocaine-dependent subjects. Nature 386:830–833.

Volkow ND, Wang GJ, Fowler JS, Logan J, Hitzemann R, Ding YS, Pappas N, Shea C, Piscani K (1996) Decreases in dopamine receptors but not in dopamine transporters in alcoholics. Alcohol Clin Exp Res 20:1594–1598.

Volkow ND, Wang G-J, Tomasi D, Baler RD (2013) Unbalanced neuronal circuits in addiction. Curr Opin Neurobiol 23:639–648.

Waaktaar T, Kan K-J, Torgersen S (2018) The genetic and environmental architecture of substance use development from early adolescence into young adulthood: a longitudinal twin study of comorbidity of alcohol, tobacco and illicit drug use. Addiction 113:740– 748.

Wang GJ, Volkow ND, Fowler JS, Logan J, Abumrad NN, Hitzemann RJ, Pappas NS, Pascani K (1997) Dopamine D2 receptor availability in opiate-dependent subjects before and after naloxone-precipitated withdrawal. Neuropsychopharmacology 16:174–182.

Wang J, Huang D, Zhou Y, Yao H, Liu H, Zhai S, Wu C, Zheng Z, Zhao K, Wang Z, Yi X, Zhang S, Liu X, Liu Z, Chen K, Yu Y, Sham PC, Li MJ (2020) CAUSALdb: a database for disease/trait causal variants identified using summary statistics of genome-wide association studies. Nucleic Acids Res 48:D807–D816.

Wang K, Li M, Hakonarson H (2010) ANNOVAR: functional annotation of genetic variants from high-throughput sequencing data. Nucleic Acids Res 38:e164.

Ward LD, Kellis M (2016) HaploReg v4: systematic mining of putative causal variants, cell types, regulators and target genes for human complex traits and disease. Nucleic Acids Res 44:D877–81.

Watanabe K, Taskesen E, van Bochoven A, Posthuma D (2017) Functional mapping and annotation of genetic associations with FUMA. Nat Commun 8:1826.

Wilar G, Shinoda Y, Sasaoka T, Fukunaga K (2019) Crucial Role of Dopamine D2 Receptor Signaling in Nicotine-Induced Conditioned Place Preference. Mol Neurobiol 56:7911– 7928.

Wiltschko AB, Pettibone JR, Berke JD (2010) Opposite effects of stimulant and antipsychotic drugs on striatal fast-spiking interneurons. Neuropsychopharmacology 35:1261–1270.

Worsley Hunt R, Mathelier A, Del Peso L, Wasserman WW (2014) Improving analysis of transcription factor binding sites within ChIP-Seq data based on topological motif enrichment. BMC Genomics 15:472.

Xu Z, Liang Q, Song X, Zhang Z, Lindtner S, Li Z, Wen Y, Liu G, Guo T, Qi D, Wang M, Wang C, Li H, You Y, Wang X, Chen B, Feng H, Rubenstein JL, Yang Z (2018) SP8 and SP9 coordinately promote D2-type medium spiny neuron production by activating Six3 expression. Development 145.

Yue F et al. (2014) A comparative encyclopedia of DNA elements in the mouse genome. Nature 515:355–364.

Zeng X, Liu D, Zhao X, Chao L, Li Y, Li H, Li W, Gui L, Wu W (2019) Association of bone mineral density with lung function in a Chinese general population: the Xinxiang rural cohort study. BMC Pulm Med 19:239.

Zhang X, Kaplow IM, Wirthlin M, Park TY, Pfenning AR (2020) HALPER facilitates the identification of regulatory element orthologs across species. Bioinformatics.

Zhou J, Troyanskaya OG (2015) Predicting effects of non-coding variants with deep learning-based sequence model. Nat Methods 12:931–934.

Zillikens MC et al. (2017) Large meta-analysis of genome-wide association studies identifies five loci for lean body mass. Nat Commun 8:80.

